# Organelle Volume Facilitates Spindle Positioning During *C*. *elegans* Female Meiosis

**DOI:** 10.64898/2026.10.07.757430

**Authors:** Denisa G Lazureanu, Eric Krueger, Wei Qiu, Aspen A Kopley-Smith, Clive S Smith, Alma P Aquino, Aastha Lele, Nadiia Voloshchuk, Daniel B Cortes, Mooncheol Park, Kunihiro Uryu, Malan Silva, Eric Trautman, Michael Innerberger, Stephan Preibisch, Roman Vaxenburg, Alyssa K Stark, Christopher K. E. Bleck, Molly Reilly, Meng C. Wang, Hari Shroff, Francis J McNally

## Abstract

Cortical positioning of oocyte meiotic spindles is required for reducing chromosome number by segregating chromosomes into polar bodies. In *C. elegans*, kinesin-1 transports membrane-bound organelles inward into a central ball that is hypothesized to push the meiotic spindle outward. Using focused ion beam scanning electron microscopy, we found that the percent volume occupied by membrane-bound organelles was 28% in a mature oocyte that has completed endocytosis of the yolk precursor, vitellogenin, compared to 17% in a less mature oocyte that has not yet endocytosed vitellogenin. We therefore hypothesized that blocking endocytosis of vitellogenin would reduce the size of the central ball of organelles packed inward by kinesin-1 and limit outward movement of the spindle. Depletion of vitellogenin or proteins required for endocytosis resulted in a smaller central volume of kinesin-coated organelles and defective spindle positioning. A computer simulation of a simple exclusion model faithfully recapitulated wildtype and mutant spindle position phenotypes. These results support a volume exclusion model for positioning acentriolar meiotic spindles.

## Introduction

Oocyte meiosis is a highly asymmetric process that reduces chromosome number over two divisions to produce one large oocyte and two tiny polar bodies (1). The resulting large haploid oocyte retains the organelles and other cytoplasmic materials required for embryonic development while the tiny polar bodies remain as undeveloped cells and contain the 3/4^ths^ expelled genetic material. To properly extrude polar bodies and ensure that the maternal genome is accurately segregated, the oocyte must position the meiotic spindle at the cortex. Improper segregation results in aneuploid offspring with genetic defects that can lead to embryonic death (2). Asymmetric, cortical positioning of the meiotic spindle is a conserved feature of female meiosis in all animals. In contrast with oocyte meiotic spindles, positioning of mitotic or spermatocyte meiotic spindles is driven by conserved mechanisms mediated by astral microtubules emanating from centriole-containing centrosomes (3,4). Because the oocyte meiotic spindles of many animals including humans and *C. elegans* are acentriolar, novel spindle positioning mechanisms are needed (5).

*C. elegans* oocyte nuclei are arranged within a U-shaped syncytial gonad. Oocytes arrested in the diakinesis substage of prophase become individualized and adopt a linear organization at the turn in the U. Adult *C. elegans* hermaphrodites have 8 to 10 individualized diakinesis oocytes in their proximal gonad, each denoted with a number (−1, −2, −3, etc.) that describes their relative distance from the spermatheca. Every 23 min (6), the oocyte closest to the spermatheca (referred to as the −1 oocyte) squeezes into the spermatheca where it is fertilized (7), initiates egg shell formation (8), undergoes nuclear envelope breakdown and initiates spindle migration to the cortex (9)(10). Thus, at any fixed time point, the −1 oocyte is poised to initiate nuclear envelope breakdown and spindle migration within 23 min. Preparation for meiotic maturation in the −1 oocyte includes changes in chromosome structure (11)(12), nuclear migration away from the spermatheca and future site of fertilization (6)(13), and endocytosis of the yolk protein, vitellogenin (14). Secretion of vitellogenin from somatic tissue followed by endocytosis in oocytes is a conserved feature in animals other than placental mammals. Vitellogenin is secreted from the liver in chickens and from the intestine in *C. elegans*, then endocytosed in oocytes via a conserved receptor (14)(15). We here define endocytosis-derived vesicles in oocytes that contain vitellogenin as “yolk granules”. Early studies detected endocytosis of vitellogenin and surface expression of the vitellogenin receptor, RME-2, predominantly in the −1 to −3 oocyte (14). However later studies found very small numbers of vitellogenin-positive vesicles as early as −6 oocytes (16). These observations suggest that endocytosis of vitellogenin begins in the −6 oocyte and ends at eggshell deposition.

Meiotic spindle assembly initiates in the −1 oocyte as the nuclear envelope fenestrates, allowing entry of tubulin. During nuclear envelope breakdown and spindle assembly, the oocyte squeezes through one sphincter of the spermatheca where it is fertilized. The prometaphase zygote then squeezes through a second sphincter into the uterus in a process called ovulation. Yolk granules and maternal mitochondria move inward at the same time the spindle moves outward during ovulation so that by metaphase I, the meiotic spindle is at the cortex in a region devoid of yolk granules and mitochondria. Both inward packing of organelles and outward movement of the prometaphase spindle require kinesin heavy chain (UNC-116), kinesin light chains (KLC-1/KLC-2), and a kinesin cargo adapter (KCA-1) (9,13)(10). Cytoplasmic microtubules are hypothesized to have minus ends anchored at the oocyte cortex with plus ends facing inwards so that when kinesin travels to plus ends, it results in the inward packing of organelles (10,13). Microtubule orientation with minus ends at the cortex is supported by the presence of γ-tubulin ring complex at the cortex of earlier oocytes (17), and the premature inward packing of yolk granules in diakinesis oocytes due to expression of a constitutively active kinesin heavy chain (10). Upon activation of the anaphase promoting complex (APC), the spindle shortens, cytoplasmic dynein is recruited to the cortex and spindle poles, and the spindle rotates to a perpendicular orientation (9,18–20). In kinesin-depleted embryos, metaphase spindles are found at variable distances from the cortex. Upon initiation of spindle shortening, cytoplasmic dynein moves the spindle to the cortex in some embryos, while perfectly centered spindles often remain stuck in the middle. Embryos with meiotic spindles that remain stuck in the middle presumably develop into pentaploid embryos that are the cause of partial embryonic lethality after germline depletion of kinesin (9).

The presumed minus-ends at the cortex orientation of cytoplasmic microtubules led to the hypothesis that inward transport of yolk granules and mitochondria indirectly forces the ER-enveloped prometaphase spindle outward by volume exclusion. In support of this model, artificially attaching kinesin motor domains to mitochondria resulted in exaggerated inward packing of mitochondria and partially rescued spindle positioning in the absence of endogenous kinesin heavy chain (10). A critical consideration in the volume exclusion model is the actual percent volume occupied by yolk granules, mitochondria, and other potential kinesin cargo before and after inward packing. In a previous analysis of one single plane electron micrograph, the percent volume occupied by all organelles in a −1 oocyte was 25% and this increased to 37% in the central packed region of a metaphase meiotic embryo (21). Because yolk granules appeared to have the greatest contribution to organelle volume, and because yolk granules are generated by endocytosis of vitellogenin in the most mature oocytes (14), we hypothesized that endocytosis of vitellogenin would increase the total percent volume occupied by organelles. To test this hypothesis, we quantified the percent volume occupied by all membranous organelles before and after endocytosis of vitellogenin. To ensure the accuracy of organelle percentage in a 3D volume and to prevent sampling bias from a single plane, assessment of the entire oocyte volume is critical. Here we report that the percent volume occupied by organelles is 17% in a −7 oocyte (before endocytosis of vitellogenin) and 28% in a −1 oocyte (preparing for ovulation) determined from Focused Ion Beam Scanning Electron Microscopy (FIB-SEM) volumes of entire oocytes. We further tested the prediction that reducing endocytosis of vitellogenin would result in a smaller packed ball of organelles during ovulation that would be defective in spindle positioning.

## Results

### Organelles occupy more volume in mature oocytes compared to immature oocytes

To test the hypothesis that endocytosis of vitellogenin increases the percent volume occupied by membrane-bound organelles, we measured the percent volume occupied by organelles before and after endocytosis of vitellogenin. We first generated focused ion beam scanning electron microscope (FIB-SEM) images of two volumes from the same young adult wild-type *C. elegans* worm that had been fixed and stained with osmium, which stains unsaturated lipids (22), to highlight all membrane-bound organelles. Target volumes were identified by X-ray tomography and cut from the resin block. One volume contained the entire −1 oocyte adjacent to the spermatheca that should have endocytosed a maximal amount of vitellogenin before ovulation (Fig. 1A) as well as the entire −2 oocyte and part of the −3 oocyte. The second volume contained an entire −7 oocyte that should have endocytosed very little vitellogenin (Fig. 1B) as well as the −6 oocyte and portions of the −5 and −8 oocytes. These 6 nm resolution volumes are available at Electron Microscopy Public Image Archive (EMPIAR-14008; EMPIAR-13896). 21 section z-slices were hand-segmented (Fig. 1A; 1B, Data S1-S4) by marking organelles based on their subcellular structure and organization (Fig. 1C). Yolk granules were identified as darkly stained vesicles with a clear rim and a lipid bilayer outside the clear rim (Fig. 1Ci), as such vesicles have previously been stained with anti-vitellogenin antibodies (16). The −1 oocyte volume also contained organelles with a reticulated staining pattern (Fig. 1C ii) that also occurred on portions of yolk granules (Fig. 1Ci yellow arrowhead). We classified these as “yolk-like granules”. From our data, there is no way to distinguish whether the reticulated staining of yolk-like granules arises due to damage from freezing/fixation, lysosomal degradation of lipids, or another cellular process.

**Figure 1:**
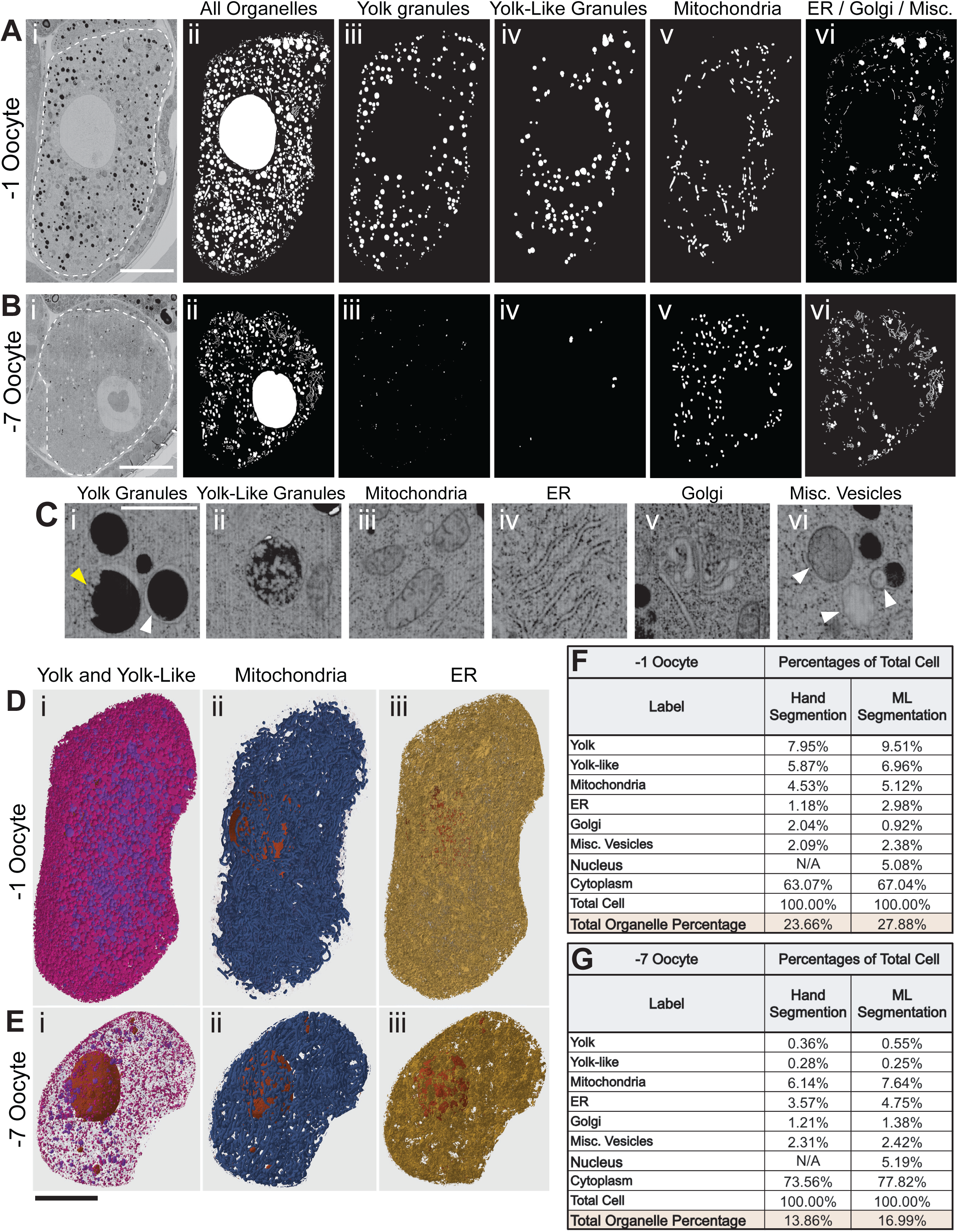
Organelles occupy more volume in mature oocytes compared to immature oocytes. (A-B) FIB Scanning Electron Microscopy image of a single focal plane of wild-type –1 (Ai) and –7 (Bi) *C. elegans* oocytes. White dashed line = plasma membrane. (ii) All the manually segmented organelles and nucleus in one focal plane labeled in white. (iii) Segmented yolk granules. (iv) Segmented yolk-like vesicles (v) Segmented mitochondria (vi) Segmented ER, Golgi, and other miscellaneous vesicles. Scale Bar = 10μm (C) Examples showing how different organelles were identified during the segmentation process. (i) Yolk granules are black with a clear rim (white arrowhead). They are mostly spherical but occasionally appear to have an irregular area with reticulated staining (yellow arrowhead). (ii) Yolk-like vesicles are mostly spherical with the majority of the vesicle having reticulated staining. (iii) Mitochondria exhibit internal cristae. (iv) Endoplasmic reticulum most often presents as stacks of thin, long lines. (v) Golgi contains closely clustered vesicles and sheets of membranes. (vi) Misc. Vesicles are vesicles that we could not distinctly identify or that occupied an extremely small volume (white arrows). Scale bar = 1μm (D-E) 3D constructions of ML segmentation of the entire −1 (D) and −7 (E) oocyte stacks. (i) Yolk (magenta) and Yolk-like vesicles (purple). (ii) Mitochondria (blue). (iii) ER (yellow). Scale bar = 10μm (F-G) Tables of percent volume data from the −1 (F) and −7 (G) oocytes shown above and calculated from hand segmentation or machine segmentation. Percentage data from second set of hand segmented −1 oocyte slices in Data S1.

To quantify the organelle classes and cytoplasmic space across the entirety of the cell, we trained machine learning (ML) models to segment the organelle classes in the −7 and −1 oocyte, respectively (Fig. 1D/1E). The training set for the −1 oocyte was from a slice hand segmented but not shown in Fig. 1A (Data S1, S3) and the −7 oocyte training set is shown in Fig. 1B (Data S1, S4). Both hand segmentation and ML segmentations are available as tiff files at DOI: 10.5061/dryad.z34tmpgx9. Values for the percent volume occupied by each non-nuclear organelle class were somewhat higher in the ML segmented full volumes than in the hand segmented representative slices (Fig. 1F, 1G). This difference may be due to sampling bias in the few slices segmented by hand. Total percent organelle volume rose from 14% in the −7 to 24% in the −1 oocyte by hand segmentation and from 17% to 28% by ML segmentation. From both data sets, the most dramatic difference between oocytes was an increase in the percent volume occupied by yolk granules + yolk-like granules which rose 20 fold in the −1 relative to the −7 oocyte (Fig. 1F, 1G; Video S1, Video S2). The total volume of the −1 oocyte was 354% of the volume of the −7 oocyte using only the ML segmented total values. 23% of this increase could be accounted for by the increase in volume of yolk granules + yolk-like granules and is thus inferred to be the result of endocytosis of vitellogenin secreted from the gut (14). If the remaining 77% of the total oocyte volume increase was due only to an increase in cytoplasmic volume, then the percent volume occupied by mitochondria would be expected to decrease from 7.64% (−7) to 2.16% (7.64/354) in the −1 oocyte. The higher observed (5.12%) volume of mitochondria in the −1 oocyte is consistent with observed transport of mitochondria into enlarging oocytes via cytoplasmic flow (23).

### Yolk granules increase in size and number across the gonad, are present past the −3 oocyte, and co-localize with KCA-1

To more rigorously connect the increasing percent volume of organelles in maturing oocytes to endocytosis of vitellogenin, we compared the number and sizes of yolk granules in representative single plane FIB-SEM images (defined as darkly staining vesicles with clear rims) (Fig. 2A-2B) with those in representative single plane confocal images of VIT-2::HaloTag (Fig. 2E). From both EM and light microscopy, the number of yolk granules was highest in the −1 and −2 oocytes but fewer yolk granules were observed as early as the −7 oocyte (Fig. 2C). The mean number values were more variable in the light microscopy because the data was from 10 different worms whereas the EM data was from multiple planes in the same worm. From both EM and light microscopy, mean sizes of yolk granules were also highest in the −1 and −2 oocytes but very small yolk granules were detected as early as the −7 oocyte (Fig. 2D). Mean sizes from FIB-SEM images were likely smaller than in confocal images because the 6 nm thick FIB-SEM images include sections through the top or bottom, rather than middle of a yolk granule and because the smaller vesicles are smaller than the resolution limit for the light microscope. These results support the idea that endocytosis of vitellogenin causes yolk granules to increase in both size and number as oocytes move from the −7 position to the −1 position.

**Figure 2:**
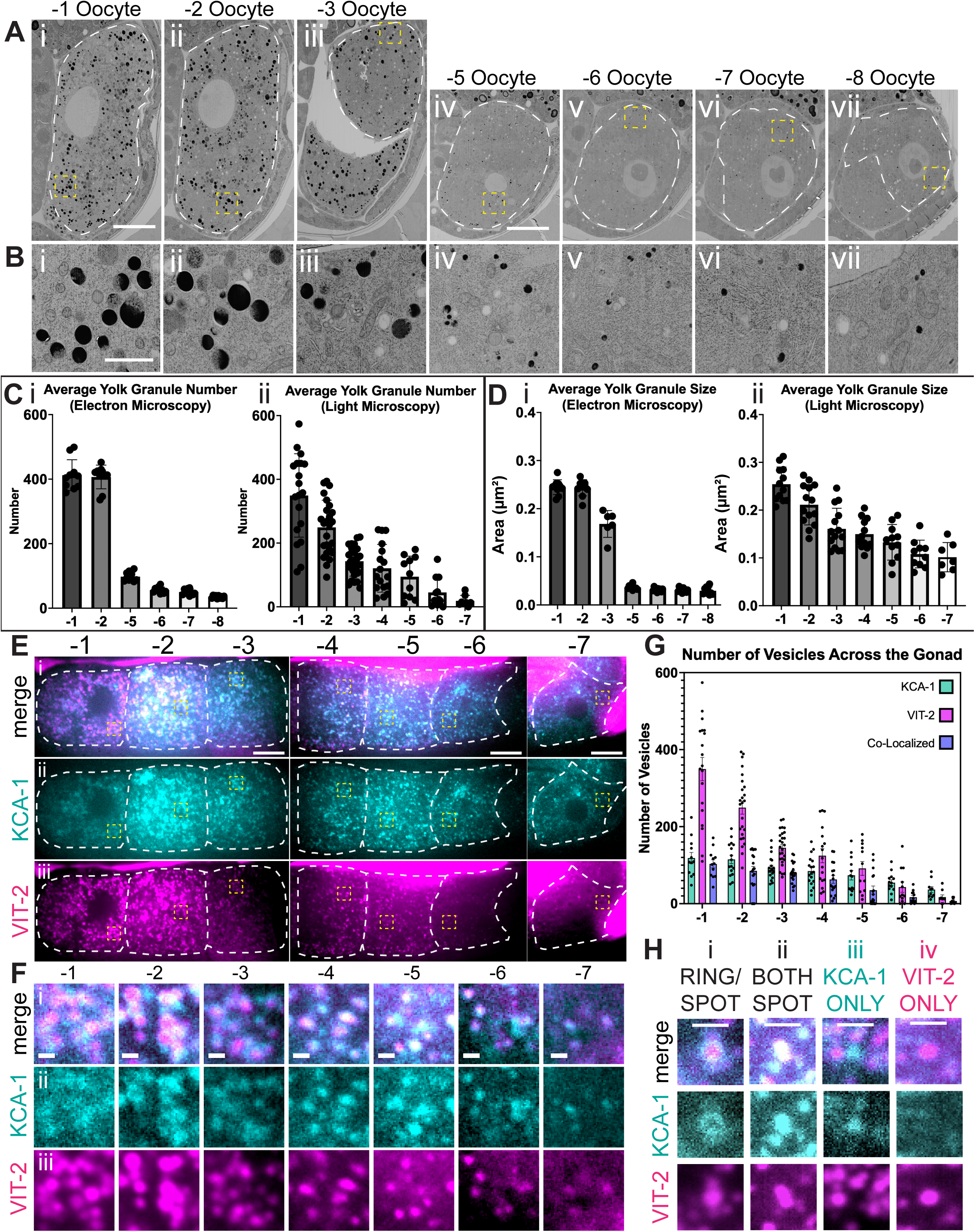
Yolk granules increase in size and number across the gonad, are present past the −3 oocyte, and co-localize with KCA-1. (A) Single plane FIB-SEM Images of the −1, −2, −3, −5, −6, −7, and −8 oocytes. White dashed line = plasma membrane. Scale bar = 10μm. Yellow Dashed Boxes = location of close-up images in Fig. 2B. (B) Close-up images of the oocyte cytoplasm corresponding to the oocytes in Fig. 2A. Black shapes are yolk granules. Scale bar = 1μm (C) Graphs showing the average number of yolk granules in each oocyte. (i) Yolk granule numbers from FIB-SEM images in each oocyte except the –3 (due to incomplete area). Each dot represents the total number of yolk granules in one focal plane, and the bar represents the average number (± SEM) from 10 focal planes. (ii) Yolk granule number from light microscopy images in each oocyte up to the −7. Each dot represents the total number of yolk granules on one focal plane, and the bar represents the average total number (± SEM) from the focal planes of different worms.(i) <u>Means ± SEM:</u> −**1**: 412.3 ± 15.15; **-2:** 406.8 ±11.64; **-5**: 97.70 ± 4.43; **-6**: 56.20 ± 2.92; **-7**: 50.30 ± 2.90; **-8**: 35.00 ± 1.35 (ii) <u>Means ± SEM:</u> −**1**: 349.6 ± 30.02; **-2:** 249.5 ±17.04; **-3**: 143.0 ±9.014; **-4**: 121.0 ±18.17; **-5**: 94.55 ± 19.43; **-6**: 45.18 ± 14.31; **-7**: 18.75 ± 6.07 (D) Graphs showing yolk granule size across the germline. (i) Data from FIB-SEM images where each dot represents the average of all yolk granule areas (µm²) in one focal plane, and the bar represents the combined average sizes (± SEM) from 10 focal planes. (ii) Data from light microscopy images where each dot represents the average of all yolk granule areas (µm²) in one focal plane in different worms. (i) <u>Means ± SEM:</u> −**1**: 0.24 ± 0.0048; **-2:** 0.24 ± 0.0053; **-3**: 0.16 ± 0.011; **-5**: 0.036 ± 0.0014; **-6**: 0.036 ± 0.0014; **-7**: 0.031 ± 0.0015; **- 8**: 0.029 ± 0.0025 (ii) <u>Means ± SEM:</u> −**1**: 0.25 ± 0.010 ; **-2:** 0.21 ± 0.011; **-3**: 0.16 ± 0.011; **-4**: 0.14 ± 0.0091; **-5**: 0.13 ± 0.011; **-6**: 0.10 ± 0.0092; **-7**: 0.10 ± 0.011 (E) Fluorescence microscopy images of a germline within a living young adult worm from the –1 to the –7 oocyte in a single focal plane and fed empty RNAi plasmid (L4440) for 24hrs. (i) Merge of fluorescent channels. (ii) Green channel (488 excitation) endogenously tagged KCA-1 with GFP. (iii) Magenta channel (640 excitation) picturing endogenously tagged VIT-2 with HaloTag and fed Janelia Fluor 646 HaloTag ligand for 24hrs. Yellow Dashed Boxes = location of close-up images in Fig. 2F. Scale bar = 10μm. (F) Close-ups of images from Fig. 2E across the germline. (i) Merge of fluorescent channels showing co-localization between KCA-1::AID::GFP and VIT-2::HaloTag. (ii) Green channel (488 excitation) endogenously tagged KCA-1. (iii) Magenta channel (640 excitation) endogenously tagged VIT-2. Scale bar = 1μm. (G) Graph of the average number of KCA-1, yolk granules, and co-localized vesicles across the oocytes. (H) Single plane examples of KCA-1 and VIT-2 localization with (i) KCA-1 in a ring around VIT-2, (ii) KCA-1 and VIT-2 co-localized as punctate, (iii) KCA-1 spot without VIT-2, and (iv) VIT-2 spot without KCA-1. Scale bar = 1μm.

We previously reported that KCA-1, the kinesin cargo adaptor that allows for the transport of yolk granules by kinesin-1, is present on the surface of vitellogenin-containing yolk granules in −1 oocytes but that KCA-1-coated vesicles were also present in earlier oocytes before endocytosis of vitellogenin (10). The existence of KCA-1-coated vesicles without vitellogenin in early oocytes suggested that KCA-1-coated “empty yolk granules” would still be kinesin cargo if endocytosis of vitellogenin were blocked. To define the relationship between vitellogenin-containing vesicles (yolk granules) and KCA-1-coated vesicles more completely, we quantified co-localization in the −1 to −7 oocytes of worms with endogenously tagged VIT-2::HaloTag and KCA-1::GFP (Fig. 2E-2H). In the −7 oocyte, VIT-2 was detected in only 19% of KCA-1 vesicles. The fraction of KCA-1 vesicles containing VIT-2 progressively increased until 90% of KCA-1 vesicles contained VIT-2 by the −1 oocyte (Fig. 2G). Whereas the number of KCA-1 vesicles and yolk granules both increased as oocytes matured, the number of yolk granules started to increase faster than the number of KCA-1 vesicles at the −4 oocyte, such that by the −1 oocyte, 75% of yolk granules did not have detectable KCA-1 (Fig. 2G). The finding that not all yolk granules in −1 to −2 oocytes have detectable KCA-1 is consistent with the previous observation that a constitutively active kinesin heavy chain/UNC-116 construct caused many but not all yolk granules to pack prematurely into a central ball in −1 and −2 oocytes (10). These results also suggest that a VIT-2 containing vesicle does not have a singular identity. Instead, it is likely that vitellogenin is continuously endocytosed until ovulation and is thus distributed among early endosomes, late endosomes, and lysosomes. Consistent with this interpretation, only 22% of yolk granules were positive for the lysosomal marker, LMP-1, in −1 oocytes (Fig. S1). Because endocytosis of vitellogenin is blocked by egg shell deposition at fertilization/ovulation and because all yolk granules are packed into a central ball by kinesin/KCA-1 at ovulation (10)(Fig. 3A, 3B), all yolk granules have likely matured into kinesin cargo by the initiation of ovulation.

**Figure 3:**
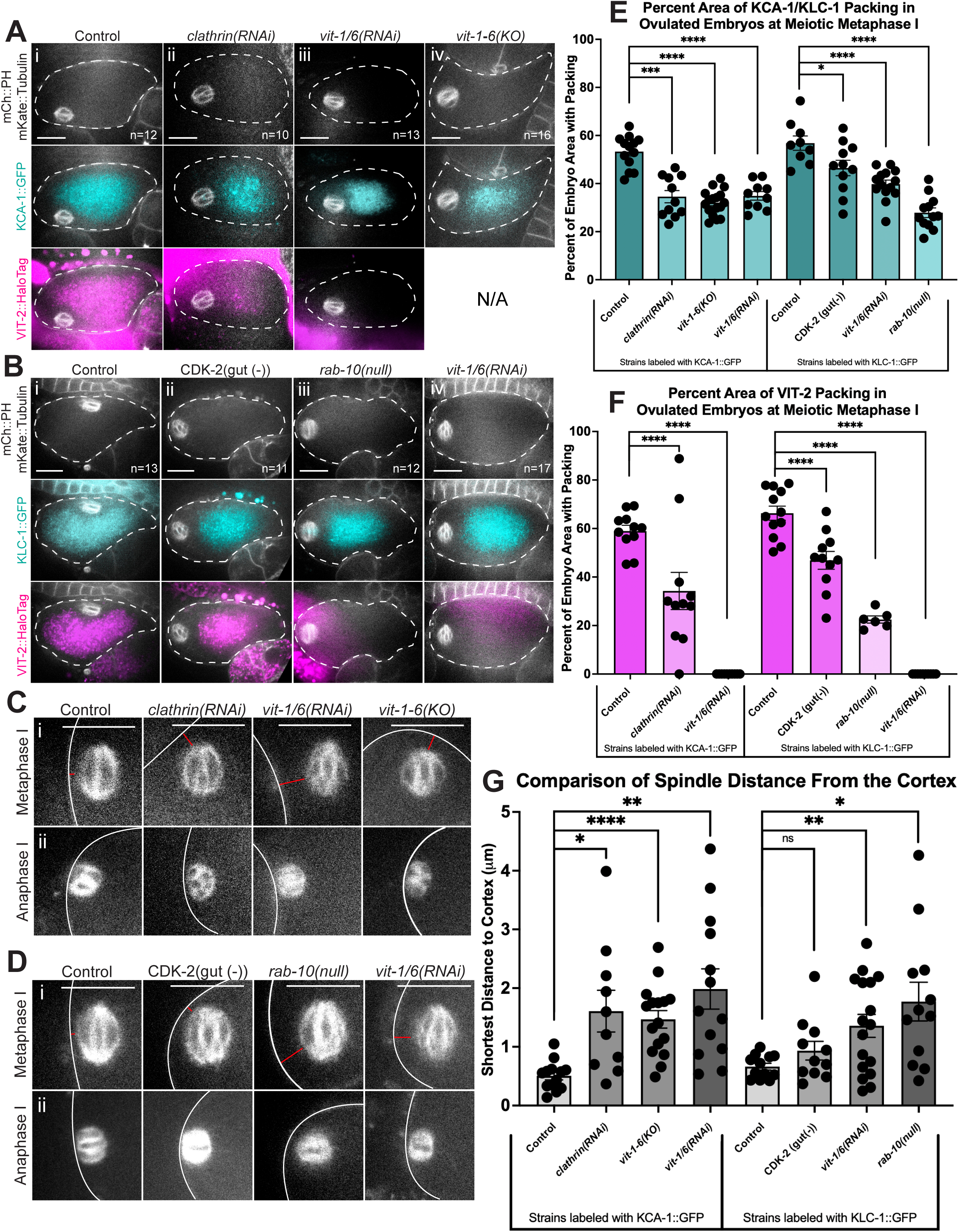
Vitellogenin Depletion Results in Tighter Organelle Packing and Spindle Localization Defects. (A) Fluorescence images taken from in utero time-lapse series of living meiotic metaphase I embryos in strains with endogenously tagged KCA-1::GFP and VIT-2::HaloTag. Tubulin and the plasma membrane are labeled with red fluorescent markers. (i) Worms fed an empty feeding clone as a control for RNAi experiments. (ii) Worms depleted of clathrin through RNAi. (iii) Worms depleted of all six vitellogenin genes through RNAi. (iv) Worm strain with knockouts in all six vitellogenin genes. Scale bars = 10μm. Dashed line drawn at the cell cortex because mCherry::PH is much dimmer than mKate::tubulin. (B) Fluorescence images taken from in utero time-lapse series of living meiotic metaphase I embryos in strains with endogenously tagged KLC-1::GFP and VIT-2::HaloTag. Tubulin and the plasma membrane are labeled with red fluorescent markers. (i) Worms fed an empty feeding clone as a control for RNAi experiments. (ii) CDK-2^gut(−)^ worms with decreased yolk production. (iii) Worms null for *rab-10*. (iv) Worms depleted of all six vitellogenin genes through RNAi. Scale bars = 10μm. Dashed line drawn at the cell cortex because mCherry::PH is much dimmer than mKate::tubulin. (C-D) Fluorescence images taken from in utero time-lapse series of living of meiotic embryos in strains with KCA-1::GFP (C) or KLC-1::GFP (D). (i) Meiotic metaphase I spindles labeled with mKate::tubulin shown in white. (ii) Meiotic anaphase I spindles from the same embryo. White line drawn at the cortex because mCherry::PH is dimmer than mKate::tubulin. Red line = shortest distance from cortex to spindle edge. Scale bar = 10μm (E-F) Graphical comparisons of the percent area of the meiotic embryo that is occupied by KCA-1/KLC-1 (E) or VIT-2 (F) across the different yolk depletion conditions. Each data point represents one worm filmed through ovulation with packing measured at meiotic metaphase I directly following exit from spermatheca. Bar represents the mean of packing percentages (± SEM). ns = not significant; * p < .05; ** p < .01; *** p < .001; **** p < .0001; Ordinary Welch ANOVA test with multiple comparisons with Dunnett correction. (E) <u>Means ± SEM:</u> **KCA-1 control**: 53.31 ± 1.87; ***clathrin(RNAi)***: 34.56 ± 2.54; ***vit-1-C(KO)*:** 32.09 ± 1.28; **KCA-1 *vit-1/C(RNAi)***: 34.86 ± 1.79; **KLC-1 control:** 56.82 ± 2.98; CDK-2^gut(−)^: 46.47 ± 3.22; **KLC-1 *vit-1/C(RNAi):*** 39.76 ± 1.70; ***rab-10(null)*:** 27.87 ± 2.01; (F): <u>Means ± SEM:</u> **KCA-1 control**: 58.92 ± 2.39, ***clathrin(RNAi)*:** 34.24 ± 7.66, **KCA-1 *vit- 1/C(RNAi)***: 0 ± 0, **KLC-1 control:** 66.32 ± 2.90, CDK-2^gut(−)^: 46.88 ± 3.67**, KLC-1 *vit-1/C(RNAi)***: 0 ± 0, ***rab-10(null)*:** 22.46 ± 1.48 (G) Graphical comparisons of shortest distance from cell cortex to spindle in worm strains with KCA-1::GFP or KLC-1::GFP. Each data point represents one worm filmed through ovulation with spindle distance measured at metaphase I. Bar represents mean of all distances (± SEM). ns = not significant; * p < .05; ** p < .01; *** p < .001; **** p < .0001; Brown-Forsythe and Welch ANOVA test with multiple comparisons with Dunnett correction. (G): <u>Means ± SEM:</u> **KCA-1 control**: 0.50 µm ± 0.068, ***clathrin(RNAi)*:** 1.60 µm ± 0.35, ***vit-1-C(KO):*** 1.46 µm ± 0.15, **KCA-1 *vit-1/C(RNAi)*:** 1.98 µm ± 0.34, **KLC-1 control**: 0.66 µm ± 0.057, CDK-2^gut(−)^: 0.93 µm ± 0.15, **KLC-1 *vit-1/C(RNAi)*:** 1.35 µm ± 0.19, ***rab-10(null)*:** 1.77µm ± 0.33.

### Blocking endocytosis of vitellogenin reduces the volume occupied by KCA-1/KLC-1 coated organelles after ovulation and reduces the efficiency of meiotic spindle translocation

The concomitant increase in size of VIT-2 containing vesicles, the increase in size of KCA-1-coated vesicles, and the increase in the percent of KCA-1 vesicles containing VIT-2 suggested that endocytosis of vitellogenin is responsible for the increasing size of KCA-1- coated endosomes. We therefore hypothesized that inhibiting endocytosis of vitellogenin would decrease the volume of KCA-1-coated endosomes, and these smaller or less numerous vesicles would pack into a smaller percent of the volume of the zygote at ovulation/fertilization. We used several genotypes previously reported to reduce endocytosis of vitellogenin and imaged endogenously tagged KCA-1::GFP or KLC-1::GFP along with VIT-2::HaloTag directly following ovulation at meiotic metaphase I. At this stage, KCA-1 and KLC-1 are still present on vesicles before switching onto filaments (10). In control worms just after ovulation, the KCA-1/KLC-1-coated organelles are packed away from the cortex such that they occupy 53 - 56% of embryo area in a mid-focal plane (Fig. 3Ai, 3Bi, 3E). There was no significant difference between the area occupied by KCA-1::GFP and VIT-2::HaloTag or between KLC-1::GFP and VIT-2::HaloTag within the same control strain (Fig. 3E, 3F). A reduced fluorescence intensity of VIT-2::HaloTag vesicles was observed after inhibiting endocytosis with *clathrin (RNAi)* (Fig. 3Aii; Fig. S2) or inhibiting vitellogenin synthesis in the gut with CDK-2^gut(−)^(24)(Fig. 3Bii). Both genetic manipulations resulted in a smaller packed area of VIT-2::HaloTag vesicles (Fig. 3F) and a smaller packed area of KCA-1/KLC-1 vesicles (Fig. 3E). *rab-10(null)* strains are defective in secretion of vitellogenin from the gut (25)(26). VIT-2::HaloTag fluorescence was not detected in 6/12 *rab-10(null)*; KLC-1::GFP; VIT-2::HaloTag embryos (Fig. 3Biii) and the remaining 6/12 embryos exhibited VIT-2::HaloTag vesicles occupying a reduced area (Fig. 3F). 12/12 *rab-10(null)* embryos had a reduced area of KLC-1::GFP vesicles (Fig. 3E). Of the 6 vitellogenin genes encoded in the *C. elegans* genome, VIT-1 to VIT-5 exhibit a high degree of sequence identity while VIT-6 is more divergent. We therefore used *vit-1/6(RNAi)* to target all 6 vitellogenins. VIT-2::HaloTag fluorescence was not detected in *vit-1/6(RNAi)* embryos (Fig. 3Aiii, 3Biv) and the area occupied by KCA-1::GFP vesicles or KLC-1 vesicles was reduced (Fig. 3A, 3B, 3E). We also found a reduced area of KCA-1::GFP vesicles in a strain with deletion or nonsense alleles in all 6 vitellogenin genes, *vit-1-6(KO)* (27)(Fig. 3E). These results show that KCA-1/KLC-1 can identify and pack vesicles even when they do not contain vitellogenin. The tighter packing seen in Figures 3A/3B and quantified in Figure 3E suggests that reduced volume of individual vesicles allows them to pack closer together.

To test the hypothesis that a reduced volume of inwardly packed organelles will reduce the efficiency of outward spindle movement, we measured the closest distance between the mKate::tubulin-labeled metaphase I spindle and the mCherry::PH-labeled cortex in strains with reduced areas of KCA-1/KLC-1-coated vesicles (see materials and methods). All the genetic manipulations except for CDK-2^gut(−)^ caused a significant increase in the distance of the metaphase spindle from the cortex (Fig. 3C, 3D, 3G). CDK-2^gut(−)^ had the weakest effect on the packed area of organelles (Fig. 3E, 3F). All spindles moved the remaining distance to the cortex during spindle shortening as previously described for a partial loss of function unc-116/kinesin heavy chain allele (19). These results support the hypothesis that the volume occupied by inwardly packed organelles is important for outward spindle migration.

Because yolk granules + yolk-like granules (16.5% volume) and mitochondria (5.1% volume) occupied the highest percent volume relative to other organelles in the −1 oocyte (Fig. 1F), we hypothesized that embryos that pack only one of these organelles would have both a smaller centrally packed ball of organelles and a spindle positioning defect (Fig. 4A- 4H). We previously reported that a chimaeric kinesin that can only transport mitochondria, TOMM-20::SSPB/ K420::iLID, can partially rescue the spindle positioning defect of a germline null allele of *unc-116*, *unc-116(gk5722) duIs1* (10). This mitochondria-only kinesin packed mitochondria into a smaller central volume than controls (Fig. 4B iv; 4H), did not pack yolk granules inward (Fig. 4C iv) (10), and had a spindle positioning defect (Fig. 4F). A constitutively active kinesin with a wild-type cargo-binding tail, K420::mKate::hinge tail, was previously found to pack yolk granules but not mitochondria during ovulation (10)(Fig. 4B ii, 4C ii) and to partially rescue spindle position in the *unc-116(gk5722) duIs1* background (10). This construct packed yolk granules into a smaller central ball than controls (Fig. 4C ii, 4G) and had a partial spindle positioning defect (Fig. 4F). The mitochondria-only kinesin formed the smallest packed ball of organelles (Fig. 4G, 4H) and had the strongest spindle positioning defect (Fig. 4F). These results support the idea that the volume of individual organelles determines the volume of the centrally packed ball of organelles and that a minimum volume of the central ball is needed to push the spindle all the way to the cortex by metaphase I (Fig. 4E).

**Figure 4:**
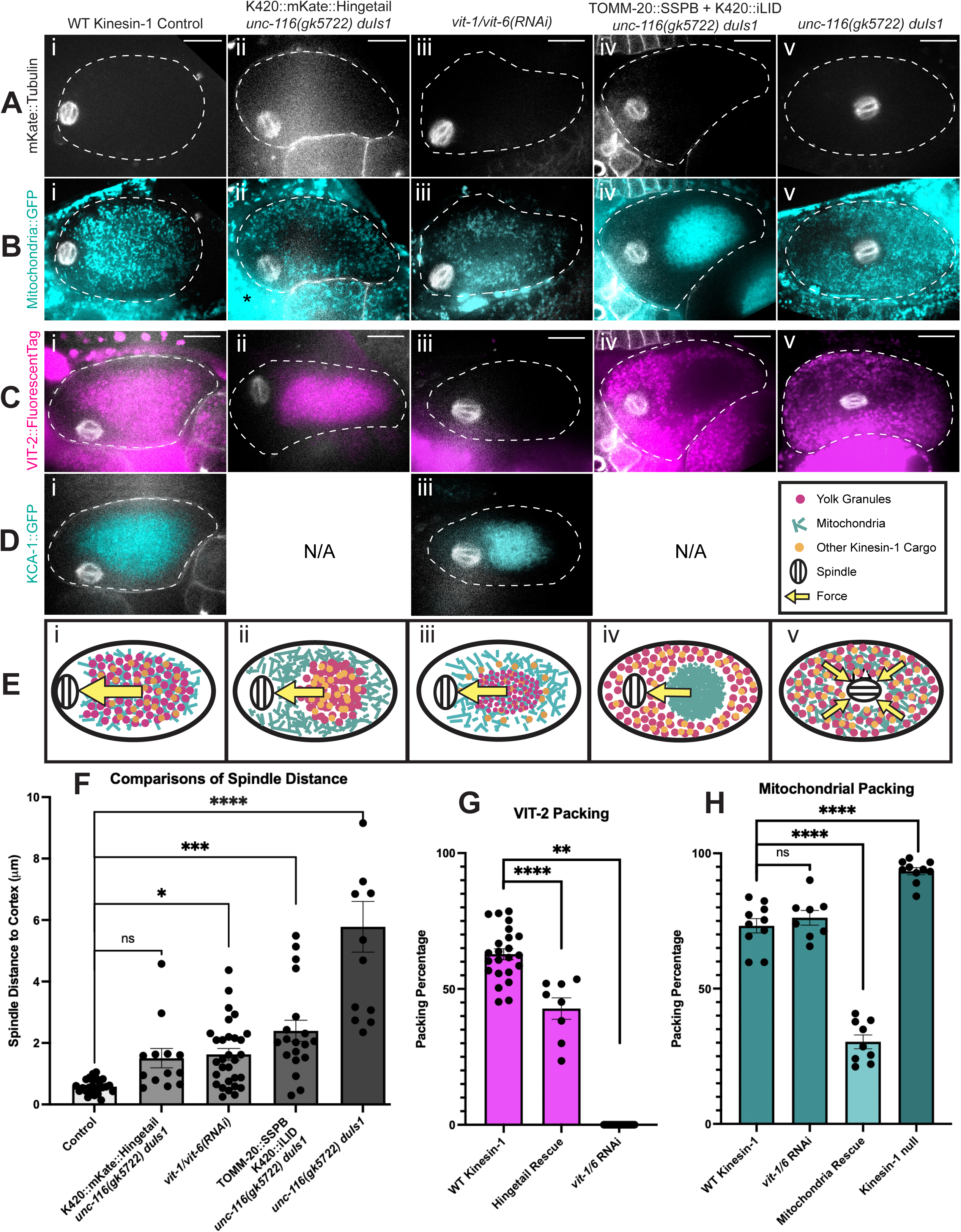
Organelle Packing Conditions and Their Effect on Spindle Localization. (A-D) Fluorescence images taken from in utero time-lapse series of living meiotic metaphase I embryos. mKate::Tubulin and mCherry::PH (plasma membrane) labeled in white (A), mitochondria (COX-4::GFP or TOMM-20::GFP::SSPB) in cyan (B), VIT-2 (VIT- 2::HaloTag or VIT-2::GFP) in magenta (C), and KCA-1 in cyan (D). (i) WT kinesin-1 condition. (ii) K420::mKate::hinge tail constitutively active kinesin-1 transgene in a kinesin-1 germline null background. (iii) Worms fed *vit-1/c* RNAi feeding clone. (iv) Optogenetic pairing of TOMM-20::GFP::SSPB (mitochondria) and K420::iLID (tailless kinesin-1) in a kinesin-1 germline null background. (v) Kinesin-1 germline null (*unc-11c(gk5722) duIs1)* condition. White dashed line = cell cortex. Scale Bar = 10μm. * = area outside embryo of bright mitochondrial autofluorescence. (E) Diagrams demonstrating the different packing conditions from Fig 4 (A-D). (i) WT packing and spindle positioning. (ii) Hinge tail packing all kinesin-1 cargos but excluding mitochondria. (iii) Vitellogenin depletion resulting in smaller endosomes but normal packing for other organelles. (iv) Optogenetic pairing of mitochondria and kinesin-1 in a kinesin-1 germline null background packs mitochondria only and excludes all other cargo. (v) Kinesin-1 germline null with no packing. Magenta dots = vitellogenin containing endosomes. Cyan lines = mitochondria. Orange dots = other kinesin-1 cargo. Yellow arrow = pushing force. (F) Graphical comparison of shortest distance from cell cortex to spindle. Each data point represents one worm filmed through ovulation with spindle distance measured at metaphase I. Bar represents mean of all distances (± SEM). ns = not significant; * p < .05; ** p < .01; *** p < .001; **** p < .0001; One-way ANOVA multiple comparisons to mean with Dunnett correction. (F) <u>Means ± SEM</u>: **control**: 0.58µm ± 0.047, **hinge tail rescue**: 1.50µm ± 0.31, ***vit- 1/C(RNAi)***: 1.62µm ± 0.19, **mitochondria rescue**: 2.39µm ± 0.34, **kinesin-1 germline null**: 5.78µm ± 0.82, (G-H) Graphical comparisons of the percent area of the meiotic embryo occupied by VIT-2 (G) or mitochondria (H) across the different packing conditions. Each data point represents one worm filmed through ovulation with organelle distribution measured at meiotic metaphase I directly following exit from spermatheca. Bar represents the mean of packing percentages (± SEM). ns = not significant; * p < .05; ** p < .01; *** p < .001; **** p < .0001; One-way ANOVA multiple comparisons to mean with Dunnett correction. (G) <u>Means ± SEM</u>: **WT**: 62.78% ± 2.01, **hinge tail rescue**: 42.78% ± 3.94, ***vit-1/C(RNAi)***: 0% ± 0 (H) <u>Means ± SEM</u>: **WT**: 73.23% ± 2.65, ***vit-1/C(RNAi)***: 76.18% ± 2.74, **mitochondria rescue**: 30.34% ± 2.54, **kinesin-1 germline null**: 93.40% ± 1.30.

### Reduced endocytosis of vitellogenin causes premature packing of kinesin-coated endosomes

Genetic manipulations that reduced endocytosis of vitellogenin by oocytes had the unexpected phenotype of premature packing of kinesin-coated endosomes in −1 and −2 oocytes (Fig. S3). Because ovulation occurs every 23 min (6), a minority (10%) of control worms show partial packing of yolk granules in −1 oocytes presumably because those oocytes are preparing for ovulation. In contrast, the majority of *clathrin(RNAi), vit-1/6 (RNAi),* or *vit-1-6(KO)* worms exhibited premature packing of KCA-1 coated vesicles in −1 and −2 oocytes (Fig. S3A). Likewise, CDK-2^gut(−)^, *rab-10(null)*, and *vit-1/6(RNAi)* worms exhibited premature packing of KLC-1-coated vesicles (Fig. S3C). In cases where VIT-2::HaloTag was reduced but still detectable, the remaining VIT-2 was also in these prematurely packing vesicles (Fig. S3A, S3C). A possible explanation for this result is that a fixed number of kinesin motors redistributes on a reduced number of endosomes such that individual endosomes acquire more kinesin motors. This increased number of kinesin motors on a subset of endosomes is sufficient to overcome the negative regulation that prevents premature packing in wild type. In addition, oocytes with reduced vitellogenin had large round volumes that excluded KCA-1 (Fig. S3B) or KLC-1 (Fig. S3D) vesicles.

### Modeling of packing and volume data demonstrates the feasibility of volume exclusion as a spindle translocation mechanism

Because the genetic manipulations that reduced endocytosis of vitellogenin had additional phenotypes that might indirectly affect meiotic spindle movements, we used 2D agent-based modeling to test the plausibility of volume exclusion as a force producing mechanism with the measured volumes of organelles from FIB-SEM data (Fig. 1). Cytosim (28) models cytoskeletal dynamics by setting the parameters of solid objects, filaments, and hands that can bind and move along filaments. Volume exclusion was emulated in this 2D space by setting steric hindrance parameters to all solid objects in the simulation. This ensured that no two objects can occupy the same coordinates. In this system, the “embryo” space was filled with “non-cargo” vesicles (grey in Fig. 5; representing the volume occupied by ER) and “cargo” with “kinesin-1” hands on its surface (yolk + yolk-like granules = magenta; mitochondria = cyan; miscellaneous vesicles = orange in Fig. 5). Additionally, the embryo was populated with random nucleating points along the cortex that produced microtubule filaments with their minus ends at the cortex and their plus ends oriented towards the center of the space. A solid object that represents the spindle started perfectly in the center of the cell to simulate the most extreme condition and both cargo and non-cargo were randomly distributed at the start of each simulation. The kinesin hands were programmed to bind a filament when in proximity and transport its cargo all the way to the plus-end where it would subsequently release from the fiber.

**Figure 5:**
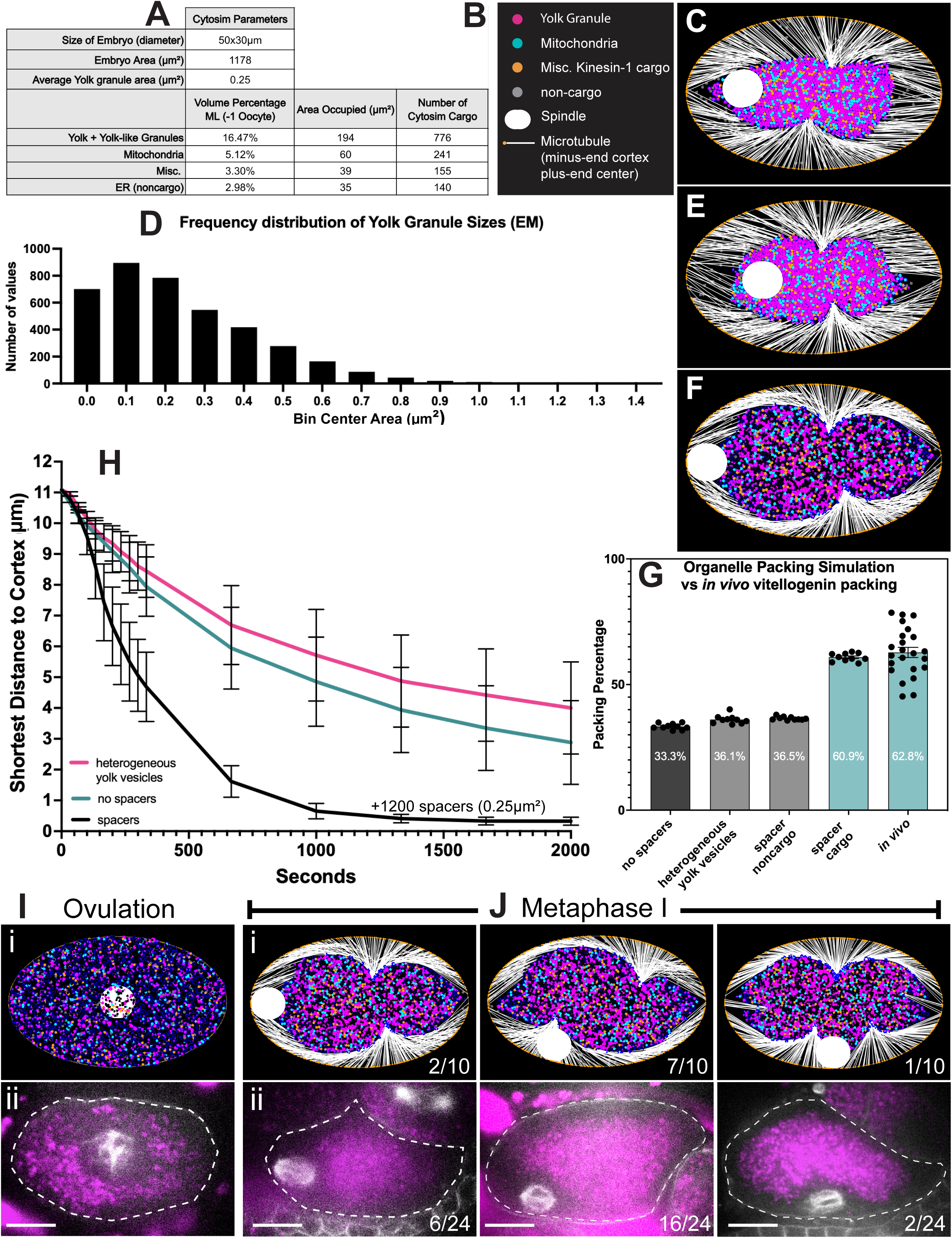
Cytosim Simulations of Organelle Packing and Spindle Movement. (A) Calculations to find the number of organelles in Cytosim modeling to match volume data from ML −1 oocyte segmentation. (B) Legend for Cytosim images. Magenta circles: vitellogenin vesicles. Cyan circles: mitochondria. Orange circles: misc. kinesin-1 cargo. Gray circles: non cargo (ER). White capsule: spindle. White lines anchored at cortex: microtubules (minus-end with orange circles at the cortex and plus-ends facing inwards). (C) Final frame at 2000s of simulation with the organelle cargo numbers calculated in (A). (D) Frequency distribution graph of yolk granule sizes from hand segmented data of the −1 oocyte (Fig. 1). (E) Final frame at 2000s of simulation after adding heterogeneous yolk granule sizes determined in (D). (F) Final frame at 2000s of simulation after adding spacer cargo (black). (G) Graphical comparisons of organelle packing percentage of the total cell in the different model iterations (10 runs) and the experimental condition. (G) <u>Means ± SEM</u>: **no spacers**: 33.3%, **heterogeneous yolk granules:** 36.1%, **spacers noncargo**: 36.5%, **spacer cargo**: 60.9%, ***in vivo***: 62.8% (H) Graphical comparisons of spindle movement in the different iterations of Cytosim modeling with each condition run 10 times. Shortest distance from the simulated spindle to the simulated cell cortex as measured in ImageJ at time points of 0s-300s, 333s, 666s, 1000s, 1333s, 1666s, and 2000s. (1pixel = 0.067um) (I) Comparisons of metaphase I spindle positions along the long axis of the embryo in Cytosim models and live fluorescence microscopy. (i) Simulation at 0.00s that simulates the oocyte prior to ovulation when all the organelles are evenly distributed, and the spindle is in the center. (ii) Single time point of an oocyte captured during live imaging directly following ovulation when the organelles are beginning to pack and the spindle is forming in the center. Magenta vesicles = VIT-2::HaloTag; White Spindle = mkate::tubulin (J) Comparisons of metaphase I spindle positions in Cytosim models and live fluorescence microscopy after ovulation. (i) Final frame of simulations at 2000.00s when the cargo has packed and the spindle is at the cortex. The spindle reaches the cortex at different locations along the cortex (ii) Single frames of a live timelapse showing the similarity of packing to simulations and the positions/frequencies of the metaphase I spindle along the cortex. Magenta vesicles = VIT-2::HaloTag; White Spindle = mkate::tubulin

Initially, a single vesicle radius of 0.28μm was chosen, based on the mean size of yolk granules from Fig. 2D. The numbers of each vesicle class required to fill the percent area corresponding to the percent volume measurements from ML-segmented FIB-SEM (Fig. 1F) were determined (Fig. 5A) and used to populate the initial simulation which was run 10 times. Under wildtype conditions, a mature oocyte will take an average of 1000s from initiation of organelle packing to exit from the spermatheca (10). In the first simulation after 2000s, cargo vesicles were packed into the center of the embryo, but the spindle did not reach the cortex consistently (Fig. 5C). However, the percent area occupied by the central ball of VIT-2 containing yolk granules relative to the entire embryo was 33%, much less than the experimentally determined size of 63% determined from Fig. 3F (Fig. 5G). Changing yolk granules from a single uniform size to heterogeneous sizes matching the frequency distribution determined from electron microscopy (Fig. 1) while maintaining the total percent area of yolk granules (Fig. 5D) had a minimal effect on the size of the central ball of organelles (Fig. 5E, 5G). To make the simulated central ball of organelles occupy an area corresponding to experimental measurements, we added 1200 additional cargo vesicles with 0.25μm radius (shown in black in Fig. 5F) to act as spacers. This number of spacer cargo vesicles was empirically determined because it enlarged the central ball of organelles to 61% of the embryo, similar to the experimental value of 63% (Fig. 5F, 5G). In addition, after 10 simulation runs, the percent area occupied by yolk granules, mitochondria, and miscellaneous cargo within the central ball of organelles was decreased to 41 ± 0.3 % (mean ± SEM; n=10), close to the experimental value of 37% (21). Under these simulation conditions that accurately reflect experimental measurements of the volume occupied by individual organelles (Fig. 1) and the percent area occupied by the central ball of organelles relative to the entire embryo (Fig. 3), the spindle was efficiently translocated to the cortex in 10/10 simulation runs (Fig. 5F, 5H, Video S3). In addition, under these simulation conditions, the spindle’s position along the long axis of the embryo closely matched experimental observations (Fig. 5I, 5J). The molecular mechanism of the “spacers” that prevent cargo from packing into direct contact is unknown. However, we found that substituting 1200 non-cargo spacer vesicles for the 1200 cargo spacer vesicles did not recapitulate the experimentally determined area of the central ball or efficiently move the spindle to the cortex.

We then simulated rescue of the *unc-116(gk5722) duIs1* with K420::mKate::hinge tail by converting mitochondria to non-cargo (Fig. 6B), vitellogenin depletion by deleting yolk granules (Fig. 6C), and rescue of *unc-116(gk5722) duIs1* with mitochondria-only kinesin by converting yolk granules and miscellaneous cargo to non-cargo (Fig. 6D). For the hinge tail kinesin rescue and mitochondria-only kinesin rescue, the number of spacers were reduced from 1200 to 500 and 1200 to 900 since the purpose of these imaginary cargo is to prevent real cargo from touching. Running each simulation 10 times for 2000s generated central organelle balls that occupied a percent of total embryo area similar to experimental observations (Fig. 6F). These simulations also resulted in spindle translocation defects similar to those observed experimentally (Fig. 6G, 6H, VideoS4-S7).

**Figure 6:**
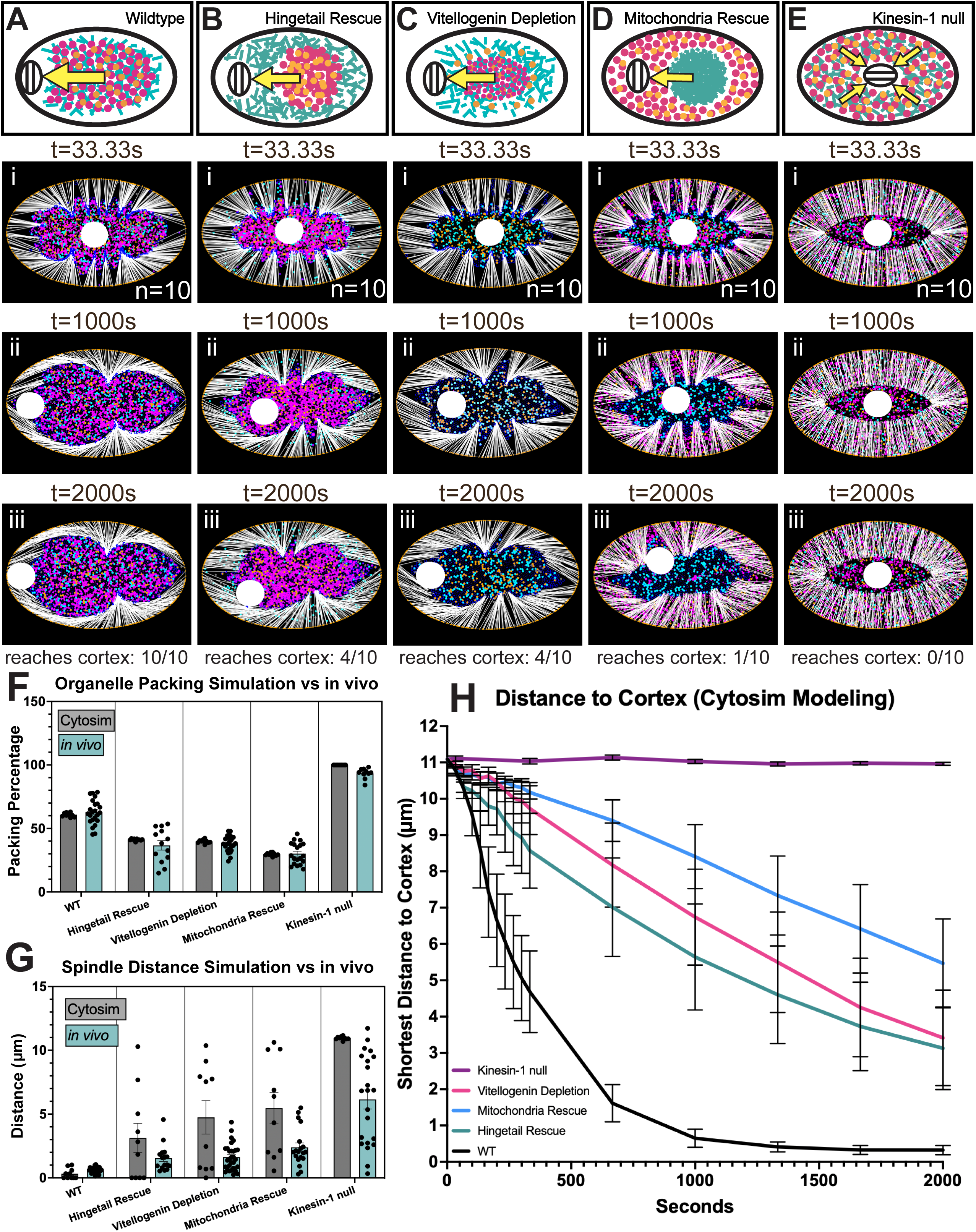
Cytosim Simulations of Different Packing Percentages Affect Spindle Localization. (A) Wild-type diagram of kinesin-1 cargo packing and spindle positioning at the cortex. (i) Frame 1 of single Cytosim simulation at 33.33s. (ii) Frame 30 of Cytosim simulation at 1000s. (iii) Frame 60 of Cytosim simulation at 2000s. Simulation run 10 times. (B) Hinge tail rescue strain (K420::mKate::hinge tail in *unc-11c(gk5722) duIs* background) diagram and simulation results. (C) Vitellogenin depletion (*vit-1/c* RNAi in a WT background) diagram and simulation results. (D) Mitochondria-only kinesin rescue (TOMM-20::SSPB and K420::iLID in *unc-11c(gk5722) duIs* background) diagram and simulation results. (E) Kinesin-1 germline null (*unc-11c(gk5722) duIs)* diagram and simulation results. (F) Graphical comparisons of organelle packing in the different simulations run in Cytosim (grey) and *in vivo* (cyan) results. (G) Graphical comparisons of final spindle distance closest to the cortex in the different simulations run in Cytosim (grey) and *in vivo* (cyan) results. (Cytosim 1pixel = 0.066801μm) (H) Graphical comparisons of spindle movement in the different conditions of Cytosim modeling with each condition run 10 times. Shortest distance from the simulated spindle to the simulated cell cortex as measured in ImageJ at time points of 0s-300s, 333s, 666s, 1000s, 1333s, 1666s, and 2000s.

## Discussion

Since the discovery of kinesin (29), models of its cellular function have primarily involved movement of cargo in the same direction as the motor, toward the plus ends of microtubules (30). Consistent with this idea, during ovulation in *C. elegans*, kinesin and its cargo (yolk granules and mitochondria) move inward from the cortex. Paradoxically, the meiotic spindle moves outward, in the opposite direction but in a kinesin-dependent manner. This led to the volume exclusion hypothesis that inward transport of direct cargo indirectly forces the meiotic spindle outward. In this model, kinesin-mediated transport of vesicles inward into a fixed space will generate an outward force once there is insufficient space for the new vesicles arriving. The plausibility of this model depends on the actual volume occupied by organelles that are packed inward. Our measurement of total organelle percent volume in a −1 oocyte (24% hand segmentation of a substack; 28% ML segmentation of the entire oocyte) was similar to a previous hand segmentation of a single thin section (25%) (21). Because yolk granules + yolk-like vesicles occupied the greatest fraction (14% hand segmentation; 16% ML segmentation), we hypothesized that blocking endocytosis of vitellogenin would reduce the volume occupied by yolk granules and yolk-like vesicles and reduce the effective outward pushing forces on the meiotic spindle. Indeed, blocking vitellogenin endocytosis [*clathrin(RNAi)*; *rab-10(null)*] or synthesis [CDK- 2^gut(−)^; *vit-1/6(RNAi)*; *vit-1-6(KO)*] reduced the volume of the large central ball occupied by organelles at metaphase I and increased the distance between metaphase I spindle and the cortex.

Analysis of FIB-SEM data of a −1 oocyte revealed 25% of oocyte volume occupied by potential kinesin cargo (yolk + yolk like+ mitochondria + miscellaneous; Fig. 1). If these vesicles fused with each other during packing, as occurs with zebrafish yolk granules (31), the giant vesicle would occupy 25% of total embryo volume. However, *C. elegans* germline yolk granules do not fuse (32) and instead, re-disperse after meiosis (13). Electron microscopy of the packed region of a meiotic metaphase embryo revealed only spherical yolk granules with space between them (21). There is a theoretical limit to how tightly uniform-sized spheres can be optimally packed into a confined space such that they can occupy a maximum of 75% of the packed volume (33). The initial Cytosim condition of uniform-sized vesicles with no spacers resulted in vesicles that occupied 74.8 ± 2.6 % (mean ± SD, n=10) of the packed area, remarkably close to the theoretical limit. However, these conditions resulted in a much smaller central ball of organelles (33% of total embryo area) than observed by light microscopy (63% of total embryo area). This discrepancy indicated that some mechanism prevents close contact of organelles during packing. Inclusion of 1200 “space filling” cargo in Cytosim mimicked this unknown mechanism and generated a central ball of organelles close to experimental values. The molecular basis of this “space filling” effect, however, is not known. One possibility is the existence of kinesin cargo that were not detected by FIB-SEM of an osmium-stained worm. The existence of these unidentified volume-displacing kinesin cargo is supported by the observation that a constitutively active kinesin, K420::mKate::hinge tail, generates a large ball that, in the absence of KCA-1, outwardly displaces mitochondria, yolk granules, and ER in −1 oocytes (10). Another mechanism that might limit inward packing include dynamic instability of the cytoplasmic microtubules and the dissociation of KCA-1 and KLC-1 from yolk granules that occurs just after ovulation (10).

Inward packing of yolk granules has been described in zebrafish oocytes (31)(34) where initially round yolk granules are forced so close together that they deform, generating flat interfaces. This has been described as forcing internal cytoplasm outward to form the blastodisc (cortical volume of cytoplasm occupied by the meiotic spindle) and a cortical yolk-free zone occupied by cortical granules. Although placental mammals do not have vitellogenins, and meiotic spindle movements are primarily actin driven rather than microtubule driven (35), there are similarities with *C. elegans* in organelle volume and organelle packing. Analysis of prophase I human oocytes, equivalent with *C. elegans* −1 oocytes, revealed total organelle occupancy of 22% (36) and a second analysis of metaphase II human oocytes revealed organelle volume occupancy of 26% (37). Mitochondria occupied an average of 4.3% - 6.3 % of total volume in human oocytes (36)(37)(38). Total organelle volume and mitochondrial volume is thus similar between human and *C. elegans* oocytes. In human oocytes, the organelles occupying the highest percent volume are a poorly characterized group of irregularly shapes vesicles called SER (smooth ER) that have been subcategorized by size (eg. SER-VLV: smooth ER very large vesicle). Total SER vesicles occupy 13% of the volume of prophase I human oocytes (36), similar to the 14% occupied by yolk + yolk-like vesicles in the *C. elegans* −1 oocyte segmented by hand (Fig. 1). In addition, both mitochondria and the abundant SER-VLV’s are packed inward away from the cortex in human prophase I oocytes (36)(38). These organelles are more evenly dispersed by metaphase II in human (37,38) and *C. elegans*. Although the exact timing of organelle packing in human oocytes has not been clearly defined, one FIB-SEM volume of a human metaphase I oocyte still had packed mitochondria and SER vesicles (38) suggesting that these vesicles may be packed during MI spindle migration. Lysosomes appear to remain packed inward in published images of human metaphase II oocytes (39). The most voluminous organelles in sheep oocytes have been referred to simply as “vesicles” and occupy 36% and 25% of pre and post maturation oocytes respectively (40). Mitochondria, lipid droplets, and other unidentified vesicles are packed inward during in vitro maturation of pig oocytes (41). Thus, organelles that occupy a much higher percent volume than mitochondria are found widely in oocytes of placental mammals that do not have vitellogenins and in some cases, these voluminous organelles pack inward during oogenesis. In mouse oocytes, mitochondria are packed inward in an actin-dependent manner before nuclear envelope breakdown and during spindle migration to the cortex (42) and it has been suggested that formin-nucleated actin polymerization from the ER enveloping the spindle pushes on this inwardly packed mass of mitochondria to drive outward spindle migration (43).

Inward organelle packing has been hypothesized to be important for a variety of other mechanisms independent of meiotic spindle migration. During *C. elegans* fertilization, inward packing initiates just before fertilization so that the sperm contents enter a volume free of yolk granules and maternal mitochondria (44). This separation of the sperm contents into a separate cytoplasmic domain may help to prevent premature interactions between maternal and paternal genomes. In addition, the layered *C. elegans* egg shell is deposited through sequential activation of chitin synthase, endocytosis of chitin synthase, and secretion of chondroitin-containing secretory vesicles. Clearing of yolk granules and maternal mitochondria from this sub-cortical zone may facilitate these regulated secretion and endocytosis events. Indeed, *kca-1(RNAi)* results in mislocalization of chondroitin-containing cortical granules away from the sub-cortex, defective secretion of these vesicles (45), and defective eggshells with compromised permeability properties (46). In mouse oocytes, a transient ball of organelles called a zollo body is packed inward in small oocytes but disperses long before meiotic maturation (47) and thus cannot play a role in spindle migration. Likewise, mitochondria and SER vesicles are dispersed by metaphase II, when fertilization occurs, in human oocytes (37)(38). Thus, inward packing of these organelles cannot be essential for fertilization-dependent events like cortical granule secretion.

The relationship between nuclear/germinal vesicle migration and early meiotic spindle positioning is variable between different species. In *C. elegans*, the germinal vesicle often migrates from a central position to the cortex at the opposite end of the oocyte from the spermatheca valve that will determine the site of fertilization (6). This nuclear migration is kinesin-dependent (13). Because the spindle is smaller than the nucleus, spindle migration still occurs when the nucleus is cortical (9). When nuclear migration does not occur, the spindle migrates over a longer distance (10). However, even when nuclear migration is blocked by kinesin depletion, the nucleus is still not perfectly centered. It has been suggested that this residual asymmetry could be generated by asymmetric growth of the oocyte (13). In contrast, the germinal vesicle is centered at nuclear envelope breakdown in the majority of mouse oocytes and the minority with a cortical germinal vesicle have a higher rate of defective meiosis resulting in aneuploidy (48). In striking contrast, the germinal vesicle migrates to a stereotypical cortical position in highly polarized oocytes like those of *Drosophila* (49), ascidians (50), starfish (51), and sea cucumbers (52). Whether there is regulated positioning of the germinal vesicle in non-rodent mammals is less clear. In multiple studies, the majority [91% (53); 70% (54); 52% (55)] of human germinal vesicle stage oocytes imaged at a single time point had a peripheral vs central germinal vesicle. However, another study tracking germinal vesical movements concluded that human germinal vesicles are extremely mobile relative to more stationary mouse germinal vesicles (56). Thus, determining the relationship between nuclear migration and spindle migration in human oocytes would require scoring the position of the nucleus precisely at germinal vesicle breakdown. Although a large number of time-lapse sequences of human oocytes starting at germinal vesicle breakdown have been reported (57), the starting position of the germinal vesicle was not reported.

## Materials and Methods

### General Maintenance of Animals

*Caenorhabditis elegans* were maintained at 20°C on MYOB plates coated with an *E. coli* OP50 lawn. Genotypes of *C. elegans* strains available in Data S6. Genotypes initially validated by Sanger sequencing of PCR products. Genotypes of strains generated through genetic crosses were validated by PCR genotyping or fluorescence microscopy. Hermaphrodites and males were used for genetic crosses to generate strains. Transgenic *C. elegans* work was carried out under a U.C. Davis Biological Use Authorization.

### RNAi Depletion and HaloTag Labeling

L3-L4 larvae were placed on IPTG-induced lawns of HT115 bacteria bearing L4440-based plasmids. *clathrin(RNAi), vit-1(RNAi)*, and *vit-6(RNAi)* acquired from Source BioScience genomic RNAi feeding library (58) and sequenced through Quintara Nanopore Sequencing to ensure matching gene sequences from WormBase (include the sequencing files??). Cultures of the feeding clones were prepared in LB+AMP (200ug/ul) and grown for 18hrs. For RNAi treatments of *clathrin* and *vit-1/vit-6*, worms were imaged live after 24hrs of feeding. Worms containing constructs with HaloTag were fed Janelia Fluor 646 HaloTag Ligand for 24hrs prior to imaging. 200μM Janelia Fluor 646 HaloTag Ligand in DMSO was diluted into M9 buffer for a final concentration of 2.5 μM. 50 μl of this dilution was pipetted onto a lawn of OP50 bacteria on a 60 mm MYOB plate with 40–80 worms.

### Live *in utero* Fluorescence Microscopy

Young adult hermaphrodites were anesthetized into a solution of 0.1% tricaine and 0.01% tetramisole in PBS on a watch glass for 30min as previously described (6,59). Worms were then transferred to a slide with a thin agarose pad (2% agarose in water). A 22-x-30-mm coverslip was placed on top and additional anesthetic was pipetted around the edges of the agarose pad under the coverslip. The slide was inverted and placed on the stage of an inverted microscope. Stage and immersion oil temperature was 21°C - 24°C. Meiotic embryos were identified by bright-field microscopy before initiating fluorescence. Exposures for Z-stack images were 200ms for all channels, and Z-stacks were captured at 0.2μm steps for oocytes in (Fig 2E-F,I Fig. S1). For all time-lapse data and single-focal plane images were acquired with a Solamere spinning disk confocal microscope equipped with an Olympus IX-70 stand, Yokogawa CSU10, Hamamatsu ORCA-Quest qCMOS (quantitative complementary metal oxide semiconductor) detector (46nm pixel size), Olympus 100x/1.35 objective, 100-mW Coherent Obis laser set at 30% power, and MicroManager software control. For time-lapse images in Fig. 3A-D and Fig. 4A-D, exposure time was 200ms for all channels. Time interval for timelapses was 5s. Focus was adjusted manually during time-lapse imaging.

### Cytosim Modeling

Cytosim software (Francois Nedelec) downloaded from github (project ID: 68278675), compiled and ran on a 2021 MAC Computer Terminal. 2D simulations of spindle positioning via volume exclusion were preferential to 3D simulations as simulating a 3D volume to the scale of a *C. elegans* zygote would require orders of magnitude more simulation time and computational resources. To approximate the effects of 3D volume exclusion, we enabled steric interactions for fibers, solids, and beads (different object classes used to simulate biological agents like microtubules, spindles, and organelles/cargo) in 2D Cytosim. Steric interactions were tuned so that only steric exclusion was considered, ensuring that cargo, spindles, and fibers, could not physically occupy the same 2D coordinates in our simulation space. Simulations were set up so that at t=0 the spindle (solid) is at the exact center of the ellipse-shaped space (approximation of the capsuloid shape of the *C. elegans* zygote) with cargo/organelles (solids) scattered uniformly throughout and microtubules (10μm) anchored cortically (with nucleators) with their plus ends pointing inward. At t=0 forces are not calculated, allowing the spindle and cargo to overlap in space (’volume’); following the addition of force calculations, solid objects exclude each other.

Configuration files were prepared in TextEditor and based upon values from area percentage data (Fig. 3E-F, Fig. 4G-H). Information about differences between simulation conditions and in-depth explanation of the wild-type configuration file can be found in Data S5. Each simulation condition was run 10 times and snapshots (.jpg) were taken at the simulated time points: 0s-300s (every 33s), 333s, 666s, 1000s, 1333s, 1666s, and 2000s. Snapshots were opened in ImageJ Software to measure shortest spindle lengths from cortex and packing percentages (1pixel = 0.067μm).

### EM Sample preparation

*C. elegans* were treated with 0.08 M trehalose in 25 mM HEPES buffer (pH 7.4) for 20 minutes. The worms were then transferred to a 6 mm gold-plated copper planchette (100 μm depth, #662, Technotrade International, Manchester, NH, USA) containing 2.5% sodium alginate in HEPES buffer, positioned on an ice-cold metal block, and aligned in a straight orientation using an eye lash tool. Immediately following polymerization with 0.2 M CaCl₂, the samples were subjected to high-pressure freezing using a Wohlwend Compact 03 system (Technotrade International, Manchester, NH, USA).

Following freezing, samples were stored in liquid nitrogen until freeze-substitution. Freeze-substitution was performed using a scheduled automated freeze-substitution system (AFS2, Leica Microsystems) in a medium containing 2% osmium tetroxide (OsO₄), 0.1% uranyl acetate, and 3% water in acetone. This was followed by infiltration with Eponate 12 resin (Ted Pella). In the final embedding step, samples were removed from the Eponate 12 resin and embedded in Durcupan ACM resin (Sigma Aldrich, set 44610) in between Aclar sheets for the subsequent FIB-SEM sample preparation.

The freeze-substitution was scheduled as follows:

-140 °C to −90 °C 3 h

-90 °C to −90 °C 24 h

-90 °C to −60 °C 6 h

-60 °C to −60 °C 12 h

-60 °C to −25 °C 7 h

-25 °C to −25 °C 12 h

-25 °C to 23 °C 12 h

### Enhanced FIB-SEM volume acquisition and reconstruction

Resin-embedded *C. elegans* samples were provided by Eric Krueger (Shroff Lab, Janelia) and Mooncheol Park (Wang Lab, Janelia) and imaged at the Enhanced FIB-SEM Imaging Shared Resource (eFIB-SR, Janelia Research Campus, HHMI). Sample blocks were mounted on copper SEM stubs with Durcupan epoxy and iteratively trimmed to the region of interest guided by µCT. Samples were sputter-coated with 10 nm gold, followed by 100 nm carbon to dissipate charge.

Two volumes (jrc_celegans-20240415 and jrc_celegans_20241007) were acquired on customized Zeiss Crossbeam-class FIB-SEM platforms (eFIB-SEM) developed at Janelia for large-volume, isotropic-resolution imaging (60, 61) (US patent 10,600,615). Both volumes were collected with identical SEM conditions: 0.9 keV landing energy, 2.0 nA beam current, InLens detector (with the energy-selective backscattered (ESB) signal also collected), around 3.1 mm working distance, 0.5 MHz scan rate, and 6 × 6 × 6 nm isotropic voxel size. Sequential block-face milling was performed with a 30 kV gallium FIB at 6 nm per slice. Both runs were executed continuously in the Janelia eFIB-SEM acquisition software, with on-the-fly drift and milling-rate corrections. jrc_celegans-20240415 comprises 5,694 slices of 12,000 × 12,833 pixels, acquired over a period of 23 days. Jrc_celegans_20241007 comprises 5,985 slices of 10,000 × 12,333 pixels, acquired over a period of 27 days.

jrc_celegans_20241007 (−1 oocyte) was processed with the Render framework (62) (github.com/saalfeldlab/render). Raw images were converted to HDF5 and imported into Render, where a per-image correction for SEM scan distortion was applied. The volume was acquired as a single tile per slice, so no in-plane montaging was required. Adjacent and nearby slices were registered using SIFT features extracted at 10% and 20% of full resolution, with correspondences filtered by RANSAC under a rigid model. Slice transforms were then solved with a distributed block optimizer (blocks of 500 slices), using affine models regularized towards rigidity. Alignment quality was reviewed by visual inspection and inter-slice cross-correlation; one slice affected by line-scan artifacts was replaced and one corrupt raw image was excluded before the final alignment. The aligned volume was exported as multi-resolution 8-bit N5 (2× downsampling per level) and converted to multi-resolution Zarr for public release and visualization.

jrc_celegans-20240415 (−7 oocyte) was registered with the eFIB-SR SIFT-based registration pipeline (SIFT_gs, github.com/gleb-shtengel/SIFT_gs), run in a Jupyter notebook with DASK parallelization. Acquisition statistics (working distance, milling voltage, field-of-view shifts) were extracted from all frames, giving an estimated slice thickness of about 6 nm. Each raw frame was flat-field corrected with a second-order polynomial background model fitted by RANSAC. SIFT keypoints (3 layers per octave, contrast threshold 0.04, edge threshold 10, σ = 1.6) were extracted from every frame and matched to the next frame with a FLANN matcher and a Lowe ratio test of 0.7. Frame-to-frame transformations were fitted as regularized affine transforms (L2 penalty of 1 × 10⁻⁵ on the scale and shear terms, shifts unregularized) by linear regression with iterative rejection of matches with residuals above 1.5 pixels. The cumulative transformations were smoothed with a Savitzky–Golay filter (101 frames, third order) to preserve image scale, and linear drift in x and y was subtracted. InLens and ESB images were fused at a 70:30 ratio, resampled with bilinear interpolation and written as a signed 16-bit MRC volume, which was converted to multi-resolution OME-Zarr at 6 nm isotropic voxels (5,694 × 13,941 × 12,451 voxels, z × y × x).

### Manual Segmentation

Manual segmentation of 21 section Z-slices was carried out with Microscopy Image Browser (60) (https://mib.helsinki.fi/).

### ML Organelle Segmentation

We trained a single multi-class 3D U-Net model (61) for each of the −1 oocyte and −7 oocyte datasets. The 3D U-Nets consisted of four levels with (isotropic) downsampling factors 2, 2, 3 between them. 32 feature maps at the first encoder level were multiplied by a factor of 3 at each downsampling step, and reduced by the same factor along the decoder, with the final decoder level producing 64 feature maps. The individual convolution blocks consisted of two conv layers, with kernel size (3, 3, 3), valid padding, each followed by a ReLU, and residual connections. We used strided convolution for downsampling and transposed convolution for upsampling. A kernel (1, 1, 1) convolution was applied to the 64 feature map output to produce the 9 (−1 oocyte) and 10 (−7 oocyte) class logits. The raw input shape was (196, 196, 196) and the output (92, 92, 92). The ground truth labels contained only 21 z-slices and we masked out the output pixels outside of these slices. We used cross-entropy loss in combination with logit adjustment (62) to handle the organelle class imbalance. We trained the models using the Adam optimizer for ∼2.5M iterations with learning rate 2.5e-5 and batch size 1. Final per-voxel class labels were taken as the argmax over the predicted class logits. We implemented the models in PyTorch (63), used gunpowder (https://github.com/funkelab/gunpowder) for data augmentation (3D rotations, axis flips, elastic deformations, intensity transformations) and data loading, and daisy (https://github.com/funkelab/daisy) for distributed inference. Inference was restricted to the cells of interest using a binary mask manually delineated in Amira (version 2021.1; Thermo Fisher Scientific, Waltham, MA, USA). Because the −1 oocyte nucleus was not present in the training data, we segmented the nucleus manually in Amira and incorporated it into the predicted −1 oocyte segmentation as a separate label.

### Quantification and Graphing

Microscopy Image Browser (MIB) in MATLAB was used to segment 3 sets of 21 FIB-SEM image Z-stacks by hand (Fig. 1A-B, Data S2). MIB software acquired volume data of each segmented organelle (Data S2). 3D reconstructions of machine segmentation in Figure 1D/1E done in MIB. Zarr files of the segmentation were downloaded from neuroglancer links as tif files and imported into MIB as model files to visualize the 3D volumes.

Fiji/Image J was used for measuring vesicle size/numbers, co-localization, spindle distance, and packing percentages. Vesicle size of yolk granules (electron microscopy) and KCA-1-coated vesicles (light microscopy) was measured by using the oval tool in Fiji to trace the outline of organelles and retrieve area. Vesicle counting and co-localization analysis was conducted in Fiji using the multi-point tool. To quantify co-localization of KCA-1 and VIT-2, a point was placed on every KCA-1 vesicle in the green channel and then checked for VIT-2 co-localization in the deep red channel. Spindle positioning distances from in vivo timelapse images were measured by first making an ROI box around the embryo cell cortex (mCh::PH) closest to the spindle and using the auto brightness/contrast feature to enhance the plasma membrane and mark the inside edge. Then by using another ROI box and the auto brightness/contrast feature on the spindle (mKate::tubulin) the edge of the spindle was marked. The final distance was acquired by using the line tool in Fiji and making a line from the marked edge of the spindle to the marked point on the cortex.

Quantifications were done in 2D on a single plane since 3D time-lapses were not acquired. To account for variance associated with taking 2D distance measurements on a 3D embryo, each condition has a minimum of 10 samples. Measurements of packing area percentage conducted in Fiji by using the polygon tool to trace the outline of the entire cell and then the area of organelle packing. Volume data, packing percentages, and co- localization analyzed in excel and graphed in GraphPad Prism 10 (Fig. 1D-G, Fig. 2C-D,G-H, Fig.3E-G, Fig. 4F-H, Fig. 5F). One-way (Fig. 3E–F; Fig. 4F-H) and Brown-Forsythe and Welch (Fig. 3G) ANOVA statistical tests for multiple comparisons to control with Dunnett correction was performed in GraphPad Prism 10 to calculate p values. Deconvolution of images was done by using Huygens Essential 311 (Fig. S1). Adobe Illustrator used to create diagrams and prepare figures.

## Acknowledgements

We thank Wyatt Korff (Project Teams, Janelia) for supervision and institutional oversight; Stephan Saalfeld (Janelia) for guidance on z-thickness correction; and Yurii Zubov (CellMap, Janelia) for the n5→zarr conversion tool used to generate the public volume export. We thank Jessica Feldman, Rob Dowen, and the *Caenorhabditis* Genetics Center, which is supported by the NIH Office of Research Infrastructure Programs (P40 OD010440), for strains. Thank you to Melissa C Lafrades, Samantha R Mena, and Sharan K Tamber for preparing crosses, genotyping, and testing strains. We thank Kari Price for critical reading of the manuscript.

This work was supported by the Howard Hughes Medical Institute (HHMI) and by grant R35GM136241 from the National Institute of General Medical Sciences.

---

**Figure S1:**
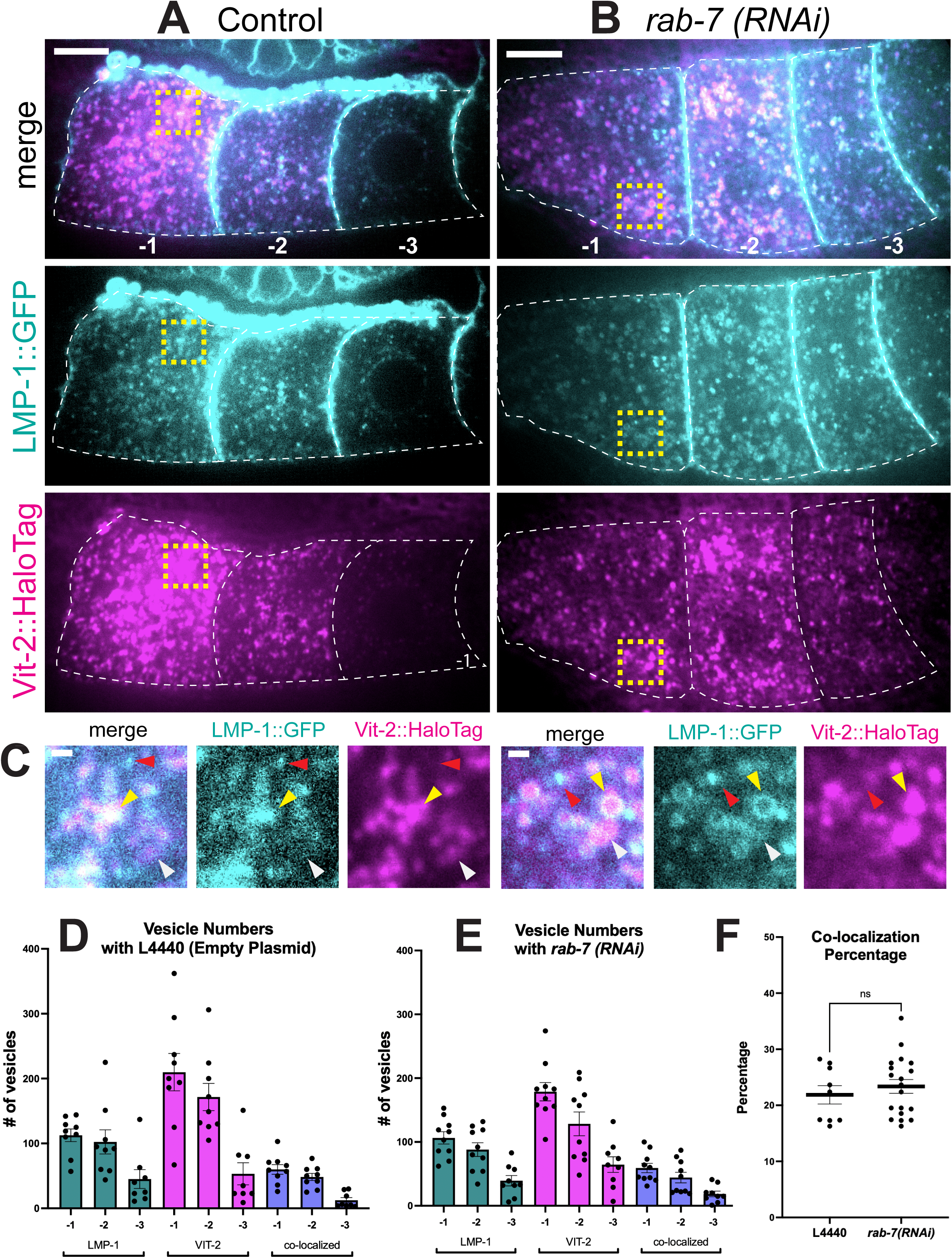
Vitellogenin co-localization with lysosomes. (A-B) Single focal plane fluorescence images of a living young adult worm germline from the –1 to the –3 oocyte. LMP-1::GFP in cyan and VIT-2::HALO in magenta. (A) Control condition where worms were fed with L4440 (empty plasmid). (B) Worms treated with *rab- 7(RNAi)* have artificially enlarged late endosomes to better visualize co-localization on the surface of vesicles. Scale Bar = 10μm (C) Enlarged images from yellow dashed squares in Fig. S1A, Fig.S1B showing vesicle localization and co-localization between lysosomal protein (LMP-1) and VIT-2. Yellow arrowhead = co-localized vesicles. Red arrowhead = LMP-1 only. White arrowhead = VIT-2 only. Scale Bar = 1μm (D-E) Graph showing the numbers of LMP-1, VIT-2, and co-localized vesicles across the germline (−1 through −3) in the control (D) and *rab-7(RNAi)* (E) condition. (F) Graph of the co-localization percentage of vesicles in both conditions. L4440: Mean: 21.87% ± 1.63; *rab-7(RNAi):* Mean: 23.36% ± 1.22

**Figure S2:**
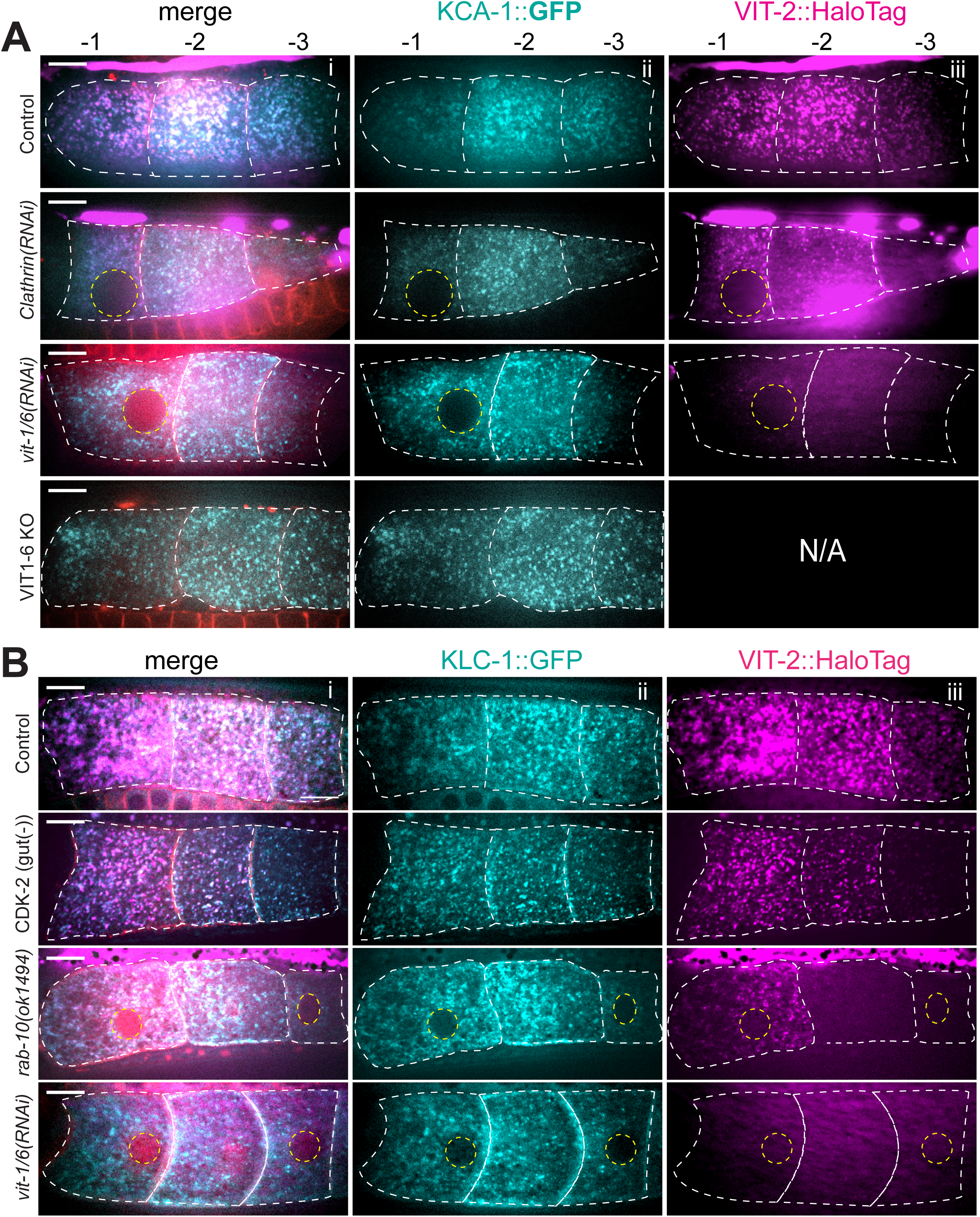
Vitellogenin Depletion Strain Gonads. (A-B) Single focal plane fluorescence images of a living young adult worm germline from the –1 to the –3 oocyte in strains and conditions with vitellogenin depletion. Some oocytes do not contain a nucleus because images are taken near the top Z-plane for better vesicle resolution. (i) Merge of fluorescence channels. (ii) Green (488 excitation) showing endogenously tagged KCA-1::GFP (A) or KLC-1::GFP (B). (iii) Magenta (640 excitation) showing endogenously tagged VIT-2::HaloTag labelled with JaneliaFluor646 HaloTag ligand. Scale bar = 10μm. White dashed line = plasma membrane of oocytes. Yellow dashed circles = nucleus outline.

**Figure S3:**
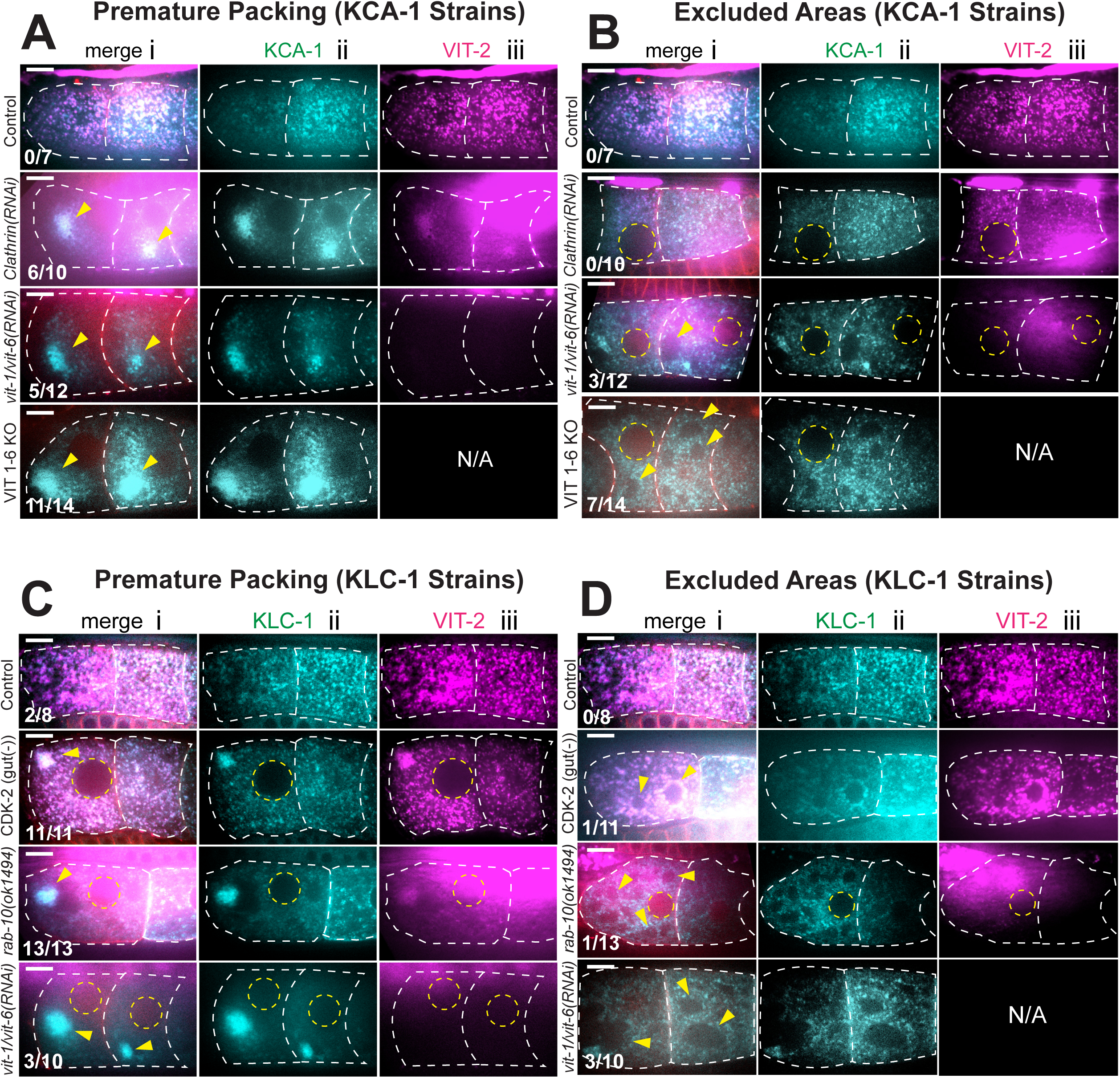
Vitellogenin Depletion Phenotypes. (A-B) Single focal plane fluorescence images of a living young adult worm germline from the –1 to the –2 oocyte in strains and conditions with vitellogenin depletion. (i) Merge of fluorescence channels. (ii) Green (488 excitation) showing endogenously tagged KCA- 1::GFP (iii) Magenta (640 excitation) showing endogenously tagged VIT-2::HaloTag labeled with JaneliaFluor646 HaloTag ligand. Scale bar = 10μm. White dashed line = plasma membrane of oocytes. Yellow dashed circles = nucleus outline. Yellow arrows: premature packing (A) and regions of exclusion (B). (C-D) Single focal fluorescence images of a living young adult worm germline from the –1 to the –2 oocyte in strains and conditions with vitellogenin depletion. (i) Merge of fluorescence channels. (ii) Green (488 excitation) showing endogenously tagged KLC- 1::GFP (iii) Magenta (640 excitation) showing endogenously tagged VIT-2::HaloTag labeled with JaneliaFluor646 HaloTag ligand. Scale bar = 10μm. White dashed line = plasma membrane of oocytes. Yellow dashed circles = nucleus outline. Yellow arrows: premature packing (C) and regions of exclusion (D).

