## Supplementary material for "Organelle Volume Facilitates Spindle Positioning During *C*. *elegans* Female Meiosis": Data S5

**Cytosim Code** 
**%%%%%%%%%%%%%%%%%%%%SIM%%%%%%%%%%%%%%%%%%%%** 
set simul system	%Sets parameters for the simulation system 
{ 
  time_step = 0.01	%(s) The interval of time that passes between each timepoint  (Brownian)
  viscosity = 1	%(µm^2^/s) Viscosity of the cytoplasm in a *C. elegans* oocyte ^1^
  steric = 1,100	%Defines whether objects/fibers can sterically repel each other

%1 turns on sterics; 100 is the energy required to violate condition

  tolerance = 0.01	%Precision of object placement
  kT = 0.0042	%(pN*μm) Thermal energy of a Brownian motion system at room temperature
}

**%%%%%%%%%%%%%%%%%%%SPACE%%%%%%%%%%%%%%%%%%%%**

set space cell	%Parameters describing a space in the simulation

{

    shape = ellipse	%Sets shape of the space

    display = ( color=0xFFFFFF66; )	%Sets display color for visualization

}

**%%%%%%%%%%%%%%%%%%%SOLIDS%%%%%%%%%%%%%%%%%%%**

set solid spindle	%Parameters for a solid object made of points

{

    steric = 1, 100	%Turns on sterics for object

    confine = all_inside, 100	%Object must stay confined inside a particular space

    display = ( style=1; color=white;)	%Sets display style and color for visualization

}

set solid yolk

{

   steric = 1, 100

   confine = all_inside, 100

   display = (color=magenta, magenta)

}

set solid mito

{

   steric = 1, 100

   confine = all_inside, 100

   display = (color=cyan, cyan)

}

set solid ocargo

{

   steric = 1, 100

   confine = all_inside, 100

   display = (color=orange, orange)

}

set solid spacer

{

   steric = 1, 100

   confine = all_inside, 100

   display = (color=black, black)

}

set solid noncargo

{

   steric = 1, 100

   confine = all_inside, 100

   display = (color=grey, grey)

}

**%%%%%%%%%%%%%%%%%%FIBERS%%%%%%%%%%%%%%%%%%%%**

set fiber microtubule	%Parameters describing a fiber in the simulation

{

   rigidity = 20	%(pN*μm^2^) approximate rigidity of microtubules ^2^

segmentation = 10	%(µm) Spacing between points along the fiber– no spacing

confine = inside, 100	%Fiber must stay confined inside a particular space

}

**%%%%%%%%%%%%%%%%%%%HANDS%%%%%%%%%%%%%%%%%%%**

set hand anchor	%Parameters describing a hand in the simulation

{

   unbinding_rate = 0	%(s^-1^) Frequency of a hand disengaging a fiber once bound

%Anchor at the cell cortex does not disengage fiber

   unbinding_force = 1e6	%(pN) Force of the force-dependent unbinding of the hand

   activity = nucleate	%Sets activity of the hand to nucleate fibers

   display = ( size = 10 )	%Sets size of hand for visualization

   nucleate = 100, microtubule, ( length=10; direction=inward; )

 %the hand will nucleate a “microtubule” with a length of 10µm with plus-ends facing inwards.

}

set single anch

{

   hand = anchor	%Defines the hand on a single agent

   stiffness = 100	%(pN/µm) Defines the deformation of the single under force

   activity = fixed	%Defines the ability of this single to move with Brownian motion

}

set hand kinesin

{

   binding_rate = 0.6	%(s^-1^) Frequency of a hand engaging a fiber

   binding_range = 0.1	%(µm) Minimal distance for a hand to bind a fiber

   unbinding_rate = 0.1	%(s^-1^) Frequency of a hand disengaging a fiber

   unbinding_force = 8	%(pN) Force-dependent unbinding of the hand

   binding_key = 1	%Key that identifies which hands can bind a fiber

   activity = move	%Sets activity of the hand to move along a microtubule

   unloaded_speed = 1	%(µm/s) Maximum translocation towards plus end with no load

   stall_force = 6	%(pN) Force-load at which motor will stall ^3^

   display = (size=2; color=blue, blue;)	% Sets display style and color for visualization

}

set single kinesin1

{

   hand = kinesin	%Defines the hand for this agent

   stiffness = 100	%(pN/µm) Defines the deformation of the single under force

}

**%%%%%%%%%%%%%%%%%%%%RUN%%%%%%%%%%%%%%%%%%%%**

new cell

{

   diameter = 50, 30	%(µm) New cell space

}

new 500 anch

{

   position = surface	%500 anchor points confined to the surface of the cell that nucleate microtubules

}

 new 1 solid spindle	%Creates a “spindle” with 3 points at an overall length of 8µm

{

point1 = 0   0  0, 3.7	%Defines a bead of radius 3.7µm at position 0

   point2 = 0.3   0  0, 3.7	%Defines a bead of radius 3.7µm at position 0.3µm from origin

   point3 = -0.3  0  0, 3.7    	%Defines a bead of radius 3.7µm at position -0.3µm from origin

   position = 0 0 0 at 0 0	%Places the spindle in the center of the cell

   orientation = 1 0 0 	%Orients the spindle horizontally

}

new 687 yolk	%687 solids that are kinesin-1 cargos

{	%Based on yolk granule size distribution (DataS1)

   sphere1= center, 0.126, 10 kinesin-1	%Defines the shape and radius (µm) of the object

%With 10 hands of kinesin on the surface

   placement= anywhere	%Placed anywhere within the cell space

}

new 293 yolk

{

   sphere1= center, 0.219, 10 kinesin-1

   placement= anywhere

}

new 154 yolk

{

   sphere1= center, 0.282, 10 kinesin-1

   placement= anywhere

}

new 77 yolk

{

   sphere1= center, 0.334, 10 kinesin-1

   placement= anywhere

}

new 46 yolk

{

   sphere1= center, 0.378, 10 kinesin-1

   placement= anywhere

}

new 25 yolk

{

   sphere1= center, 0.418, 10 kinesin-1

   placement= anywhere

}

new 12 yolk

{

   sphere1= center, 0.455, 10 kinesin-1

   placement= anywhere

}

new 6 yolk

{

   sphere1= center, 0.489, 10 kinesin-1

   placement= anywhere

}

new 3 yolk

{

   sphere1= center, 0.52, 10 kinesin-1

   placement= anywhere

}

new 1 yolk

{

   sphere1= center, 0.55, 10 kinesin-1

   placement= anywhere

}

new 1 yolk

{

   sphere1= center, 0.578, 10 kinesin-1

   placement= anywhere

}

new 241 mito

{

   sphere1= center, 0.28, 10 kinesin-1

   placement= anywhere

}

new 156 ocargo

{

   sphere1= center, 0.55, 10 kinesin-1

   placement= anywhere

}

new 140 noncargo	%140 solids that are NOT kinesin-1 cargos (do not pack)

{

   sphere1= center, 0.28	%Defines the shape and radius (µm) of the object

   placement= anywhere	%Placed anywhere within the cell space

}

new 1200 spacer

{

   sphere1= center, 0.25, 10 kinesin-1

   placement= anywhere

}

run 200000 system	%Runs the simulation

{

    nb_frames = 60	%Number of frames

}

| **Condition** | **CytoSim Conditions** |
| --- | --- |
| **WT**  **(-1)** | Cargo  1305 – “heterogeneous yolk”  156 – “other cargo” - 0.28µm radius  241 – “mitochondria” - 0.28µm radius  NonCargo  140 – “noncargo” - 0.28µm radius  Spacer Cargo (0.25µm): 1200 |
| **Hinge tail Rescue**  **(K420::mkate::hinge tail**  ***unc-116(gk5722) duIs)*** | Cargo  1305 – “heterogeneous yolk”  156 – “other cargo” - 0.28µm radius  NonCargo  140 – “noncargo” - 0.28µm radius  241 – “mitochondria” - 0.28µm radius  Spacer Cargo (0.25µm): 500 |
| ***vit-1/6* RNAi** | Cargo  0 – “heterogeneous yolk”  156 – “other cargo” - 0.28µm radius  241 – “mitochondria” - 0.28µm radius  NonCargo  140 – “noncargo” - 0.28µm radius  Spacer Cargo (0.25µm): 1200 |
| **Mitochondria Rescue**  **(TOMM-20::GFP::SSPB**  **K420::mkate::ILID**  ***unc-116(gk5722) duIs1)*** | Cargo  241 – “mitochondria” - 0.28µm radius  NonCargo  140 – “noncargo” - 0.28µm radius  1305 – “heterogeneous yolk”  156 – “other cargo” - 0.28µm radius  Spacer Cargo (0.25µm): 900 |
| **Kinesin-1 null**  (***unc-116(gk5722) duIs1)*** | Cargo  NonCargo  140 – “noncargo” - 0.28µm radius  241 – “mitochondria” - 0.28µm radius  1305 – “heterogeneous yolk”  156 – “other cargo” - 0.28µm radius  Spacer Cargo (0.25µm): 1200 |
